# Stricturing Crohn’s disease single-cell RNA sequencing reveals fibroblast heterogeneity and intercellular interactions

**DOI:** 10.1101/2023.04.03.534781

**Authors:** Pranab K. Mukherjee, Quang Tam Nguyen, Jiannan Li, Shuai Zhao, Stephen M. Christensen, Gail A. West, Jyotsna Chandra, Ilyssa O. Gordon, Sinan Lin, Jie Wang, Ren Mao, Douglas Czarnecki, Carla Rayan, Prerna Kotak, Thomas Plesec, Samir Lal, Thomas Fabre, Shoh Asano, Kathryn Bound, Kevin Hart, Chanyoung Park, Robert Martinez, Ken Dower, Thomas A. Wynn, Shaomin Hu, Nayden Naydenov, Martin Decaris, Scott Turner, Stefan D. Holubar, Scott R. Steele, Claudio Fiocchi, Andrei I. Ivanov, Kellie M. Kravarik, Florian Rieder

## Abstract

**Background:** Fibroblasts play a key role in stricture formation in Crohn’s disease (CD) but understanding it’s pathogenesis requires a systems-level investigation to uncover new treatment targets. We studied full thickness CD tissues to characterize fibroblast heterogeneity and function by generating the first single cell RNA sequencing (scRNAseq) atlas of strictured bowel and providing proof of principle for therapeutic target validation.

**Methods:** We performed scRNAseq of 13 fresh full thickness CD resections containing non-involved, inflamed non-strictured, and strictured segments as well as 7 normal non-CD bowel segments. Each segment was separated into mucosa/submucosa or muscularis propria and analyzed separately for a total of 99 tissue samples and 409,001 cells. We validated cadherin-11 (CDH11) as a potential therapeutic target by using whole tissues, isolated intestinal cells, NanoString nCounter, next generation sequencing, proteomics and animal models.

**Results:** Our integrated dataset revealed fibroblast heterogeneity in strictured CD with the majority of stricture-selective changes detected in the mucosa/submucosa, but not the muscle layer. Cell-cell interaction modeling revealed CXCL14+ as well as MMP/WNT5A+ fibroblasts displaying a central signaling role in CD strictures. CDH11, a fibroblast cell-cell adhesion molecule, was broadly expressed and upregulated, and its pro-fibrotic function was validated by NanoString nCounter, RNA sequencing, tissue target expression, *in vitro* gain- and loss-of-function experiments, proteomics, and two animal models of experimental colitis.

**Conclusion:** A full-thickness bowel scRNAseq atlas revealed previously unrecognized fibroblast heterogeneity and interactions in CD strictures and CDH11 was validated as a potential therapeutic target. These results provide a new resource for a better understanding of CD stricture formation and opens potential therapeutic developments.

## INTRODUCTION

Inflammatory bowel diseases (IBD) are a group of chronic inflammatory disorders that include Crohn’s disease (CD) and ulcerative colitis (UC).^1^ A major complication of CD is intestinal fibrosis leading to stricture formation, the main cause for surgical interventions.^2^ Conventional anti-inflammatory therapies can only modestly reduce the occurrence of stricturing disease^3, 4^ and no anti-fibrotic or selective anti-stricture therapy is available, making this a significant clinical problem.

Mesenchymal cells, the major effector cell in intestinal fibrosis,^5^ represent a distinctive cell population where fibroblasts are the predominant cell type. In the strictured bowel of CD patients, fibroblasts become activated, increase in number, and produce excessive extracellular matrix (ECM). In addition, under inflammatory conditions like CD, mesenchymal cells interact intimately with each other as well as with nearby immune and non-immune cells,^5^ leading to complex fibrogenic circuits. Until now the understanding of such circuits has been hindered by the traditional classification of mesenchymal cells into fibroblasts, myofibroblasts and smooth muscle cells,^6^ which lacks specificity.

A better understanding of the multifaceted cellular networks underlying fibrogenesis and stricture formation in CD requires high-resolution molecular profiling, such as single cell RNA sequencing (scRNAseq).^5^ This approach has revealed distinct changes in the epithelial, immune and stromal cell compartments in both in UC^7, 8^ and CD.^9^ However, as innovative as it is, the obtained information was derived from superficial endoscopic mucosal biopsies which, given the transmural nature of CD, fail to comprehensively capture the complexity of intestinal fibrogenesis which can only be fully appreciated by studying it in all intestinal tissue layers. Therefore, to comprehensively examine the transmural changes in CD and differentiate possible cellular sub-types in strictured (CDs), non-strictured (CDi) and non-involved (CDni) tissues, we leveraged scRNAseq and transcriptomic signatures in a well phenotyped cohort of CD and control surgically resected intestinal tissues.

Our approach identified previously unrecognized mesenchymal cell populations, distinct cell niches and the related cell interactions in strictured CD bowel. We identified differentially expressed genes and pathways, and comprehensive *in vitro* and *in vivo* studies demonstrated that cadherin-11 (CDH11) is a fibroblast-selective surface molecule with prominent pro-fibrotic properties.

## METHODS

### Tissue procurement

Briefly, full-thickness freshly resected intestinal specimens from subjects with CD and controls, comprising of apparently healthy tissue (constipation, healthy margin of resections from intestinal cancer patients; termed ‘NL’ for normal) were procured as previously described.^10, 11^ This protocol was approved by the Cleveland Clinic Institutional Review Board (IRB number 17-1167). Prior to resection in CD patients, the presence of a stricture was ascertained by cross-sectional imaging (computed tomography or magnetic resonance imaging) and/or inability to pass an adult or pediatric ileocolonoscope. Post resection, CD specimens were classified based on gross anatomy into segments of CDs, CDi (represented purely inflammatory disease without the presence of stricture) and CDni (non-inflammatory disease without the presence of stricture). For generation of the scRNAseq atlas the three CD segments were from the same patient’s resection. Each segment was scored and characterized using a macroscopic score modified from the simple endoscopic score (SES)-CD^10^ and a histopathologic score for inflammation^11^ or fibrosis^12^ (Supplemental Tables 4 & 5). This procurement system was validated by macroscopic and histopathologic evaluation performed by a trained IBD pathologist (I.O.G.). The time from resection in the operating room to starting tissue processing in the laboratory was < 30 minutes.

### Extracellular matsrix deposition assay for human intestinal myofibroblasts

Deposition of ECM by intestinal myofibroblasts was assayed using modification of a method described previously.^13^ Briefly, recombinant human CDH11-Fc chimera (CDH11Fc, R&D Systems; Cat# 1970-CA) or H1M1 antibody (prepared from hybridoma PTA-9699 (ATCC, Manassas, VA) at high purity (>99%) and low endotoxin levels (<0.23 EU/mg)) were used as activator and as a functional blocker of CDH11 in HIMF, respectively. A 96-well dark walled imaging plate (Greiner, BioOne, cat# 655090) was coated with CDH11Fc, H1M1 or human IgG1 or IgG2b isotype control, respectively, in PBS for 2 hours at 37℃. HIMFs were plated (3,200 cells/well) in Dulbecco’s modified Eagle medium (DMEM) supplemented with 10% fetal bovine serum (FBS) and antibiotics, and incubated overnight. Cells were then incubated with serum free culture medium to produce ECM for 5 days and subsequently removed using 0.25 M ammonium hydroxide in 50 mM Tris pH 7.4. Deposited ECM was fixed by exposure to 100% methanol at −20°C and stained with Alexa Fluor488-conjugated anti-fibronectin (EBioscience, clone FN-3, 1:500 dilution) or Alexa Fluor594-conjugated (Invitrogen, 1:1000 dilution) anti-ColI/III antibodies (EMD, Millipore Corp. 1:100 dilution). Fluorescence intensities were obtained using a Cytation5 scanner (Agilent Biotek, Santa Clara, CA). The following compound inhibitors were used in the ECM deposition assay in conjunction with the HIMF: GSK690693 [pan-Akt inhibitor] (Sigma-Aldrich; Cat# 937174-76-0), mTOR Inhibitor XI, Torin1 (Sigma-Aldrich Cat# 1222998-36-8); LY-294,002 hydrochloride [Pi3K inhibitor] (Sigma-Aldrich Cat# 934389-88-5); Y15 [focal adhesion kinase (FAK) inhibitor] (1,2,4,5-benzenetetramine tetrahydrochloride) (Sigma-Aldrich Cat# 406-66-5); SEW 2871.

Full information on Tissue dissociation and isolation of single cells, library generation, single cell RNAseq analysis, cyclic immunofluorescence, quantitative reverse transcriptase polymerase chain reaction, immunostaining and quantification, immunofluorescence labeling and confocal microscopy, RNA extraction from FFPE tissue, gene expression analysis using NanoString nCounter platform, isolation of primary human intestinal cells, cell cultures, immunoblotting, proteomics, RNA interference, next generation RNA sequencing, antibodies, animals, dextran sodium-sulfate induced colitis, chronic DSS model with CDH11 blockade in BALB/c WT mice, experimental fibrosis endpoints & statistical analysis can be found in Supplementary Materials. The GEO datasets will be made publicly available upon publication of the current study.

## RESULTS

### A single cell atlas of full thickness Crohn’s disease reveals shifts in cell type composition from non-involved to inflamed to stricturing phenotype

We generated a scRNAseq transcriptional atlas from 13 CD full-thickness ileal and 7 NL tissues. CD resections were separated into CDni, CDi and CDs segments as described in *Methods*. We mechanically dissected the MP from the mucosa/submucosa, which was subsequently processed into an epithelial-enriched fraction (LP1) and a lamina propria-enriched fraction (LP2) for a total of 99 analyzed fractions. This resulted in 409,001 high-quality single cell transcriptomes (Figure 1A). 97,954 cells were sequenced from CDni, 101,164 from CDi, 94,049 from CDs and 115,834 from NL, with a median recovery of 471 genes/cell and 751 UMI/cell (Supplemental Figure 1A). Overall preparation quality was high to excellent (viability >80%, mitochondrial reads < 10% in more than 80% of cells; Supplemental Figure 1B).

**Figure 1.**
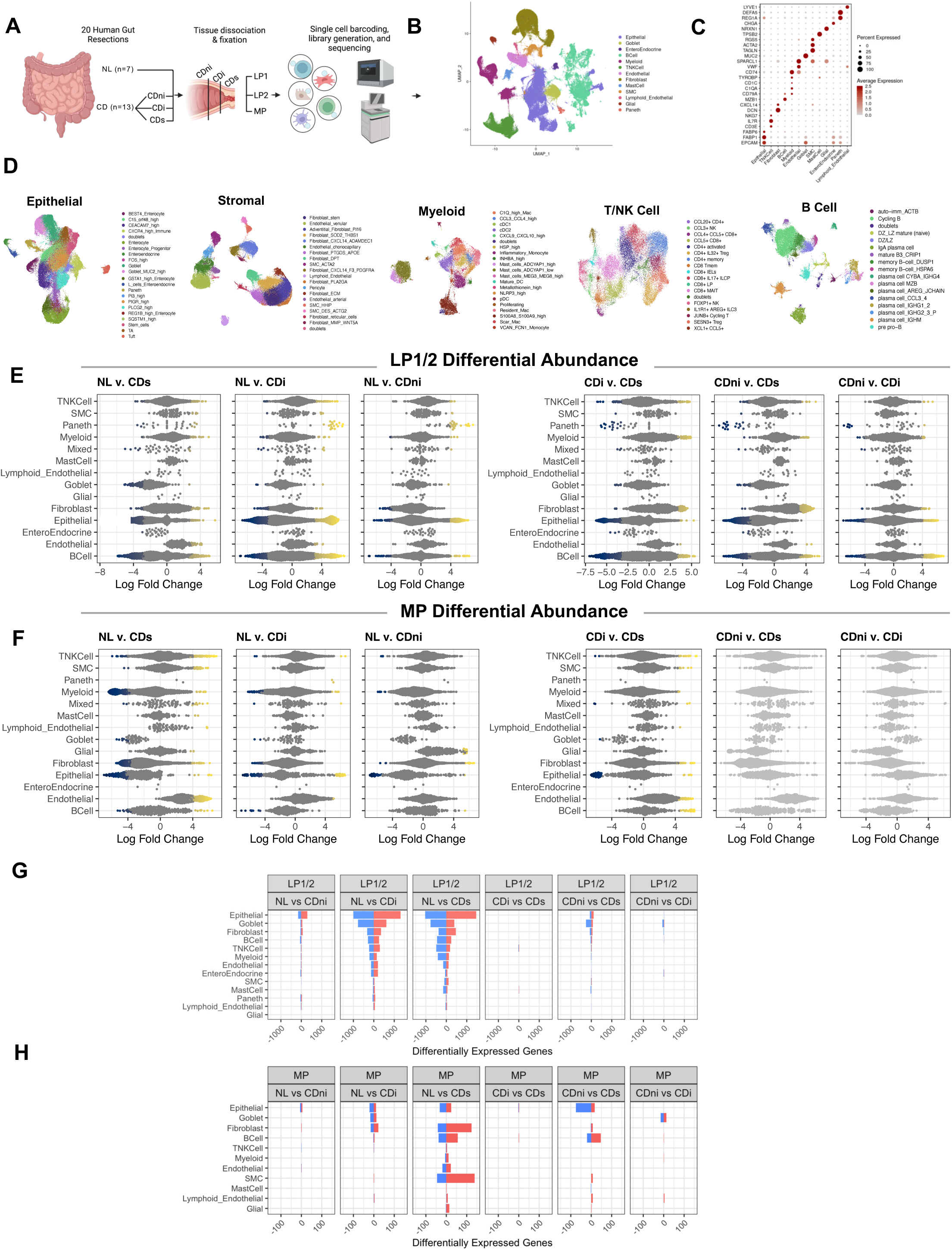
Generation of a full thickness scRNAseq atlas. **A**. Schematic summary of atlas generation from contributing donors. **B.** Uniform manifold approximation and projection (UMAP) plot of all cells from the atlas colored by cell type. **C.** Dot plot of canonical markers used to define and annotate general cell types. Average gene expression is shown by color intensity. Percent of cells expressing each gene is shown by size. **D.** Compartmental UMAP plots for each subclustering analysis colored by cell types/states. **E&F.** Differential abundance beeswarm plots from LP1/2 (**E**) or MP layer (**F**) layers by cell type. Each dot is a neighborhood of cells calculated using miloR. Neighborhoods that reach significance (spatial FDR < 0.1) are colored by log fold-change. **G&H**. Bar plot representing the total number of up-(red) or down-regulated(blue) genes by cell type in the LP1/2 (**G**) and MP (**H**) layers. Abbreviations: CD, Crohn’s disease; NL, Normal; ni, non-involved; i, inflamed, non-strictured; s, strictured.

Next, we broadly annotated the full dataset using canonical cell type markers and subclustered into epithelial, stromal, myeloid, B cell, and T/NK cell compartments (Figure 1B to D). The sequenced cell numbers per compartment and cell type can be found in Supplemental Table 6. The subclustering resulted in 93 different cell types representative of the cell populations in each segment (Figure 1C&D). To investigate progression towards strictures we examined differences in cell type composition by tissue type and layer (Figures 1E&F, Supplemental Figure 1C). Cell type composition of LP1 or LP2 versus MP was markedly different, but not between LP1 and LP2, which were thereafter combined as ‘LP1/2’. Consistent with progression from CDni to CDi, lymphocyte, myeloid, epithelial, and fibroblast cell composition was altered in CDi LP1/2 compared to CDni. Alteration in myeloid, lymphocyte, and epithelial cells was also evident in CDni compared to NL, suggesting the influence of low-grade inflammation or an intrinsic disease-induced difference in CDni (Figure 1E). In the MP layer we found significant alteration of nearly all cell types in CDs compared to NL, with minor differences in CDi and CDni compared to NL. When comparing CDs to CDi in both LP1/2 and MP, we found significant differences of epithelial, lymphocyte, smooth muscle cell, endothelial, fibroblast, and myeloid cell composition (Figure 1E&F).

In addition to cell type compositional changes related to disease progression, we examined general transcriptional changes using pseudobulk differential expression. Most differences between segments were observed in the LP1/2 layer, especially when comparing diseased to normal gut with the most marked in CDs compared to NL (Figure 1G). Epithelial cells in the LP1/2 layer exhibited the largest transcriptional changes with 2501, 2323 and 446 dysregulated genes in CDs, CDi, and CDni compared to NL, respectively. Fibroblasts also demonstrated extensive transcriptional shift with disease progression with 842, 686 and 104 dysregulated genes in CDs, CDi, and CDni compared to NL, respectively. Surprisingly, only minor transcriptional changes in the MP layer were noted between segments, even when comparing CDs to all others (Figure 1H), suggesting that major compositional changes between segments were driven by the LP1/2 layers. Related signaling pathway perturbations were also noted (Supplementary Results and Supplementary Figure 2A&B)

### Unique cell populations are represented in the stromal compartment of Crohn’s disease strictures

In the stromal compartment we identified 19 distinct cell types primarily grouped into fibroblast, smooth muscle and endothelial populations (Figure 2A&B). Among all mesenchymal cell types, only four were distinctly more abundant in the LP1/2 layer (CXCL14+ and MMP/WNT5A+ fibroblasts and HHIP/NPNT+ smooth muscle cells (SMCs), while the rest were more abundant in the MP (Supplemental Figure 1D). Interestingly, six fibroblast populations (SOD2/THBS1+, PLA2GA+, PTGDS/APOE+, MMP/WNT5A+, fibroblast reticular cells and ECM^high^) were over-represented in CDs compared to both CDi and CDni in the LP1/2 layer, particularly in CDs versus CDni. These six fibroblast populations all express markers linked to fibrotic diseases via TGF-β1 regulation, macrophage activation/differentiation, and ECM production. Among the fibroblast populations overrepresented in CDs only the MMP^+^/WNT5A^+^ fibroblasts were more abundant in any segment versus normal, specifically in CDs compared to NL (Figure 2C). Conversely, the MP layer showed minimal significant differences in cell population abundance when comparing CDs and CDi, but adventitial, stem-like, and SOD2/THBS1+ fibroblasts were under-represented and endothelial cells were over-represented in CDs compared to normal (Figure 2D). Together, these observations emphasize the LP1/2 layer as a key pathogenic area of stricturing disease as evidenced by both the global and mesenchymal compositional and transcriptional alterations in disease.

**Figure 2.**
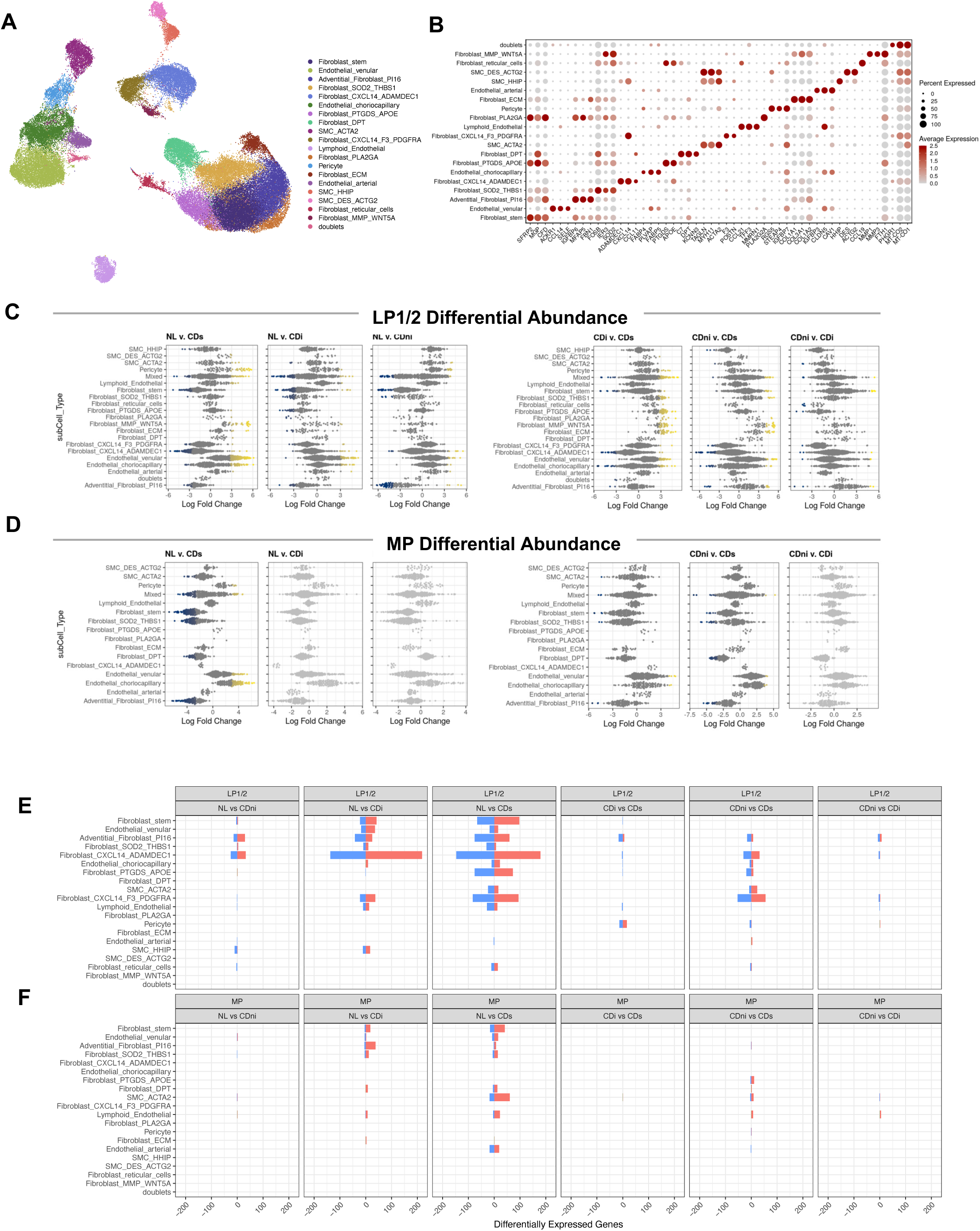
Stromal compartment analysis of the scRNAseq atlas. **A**. UMAP plot of mesenchymal subclustering colored by cell type. **B.** Dot plot of top markers used to define/annotate mesenchymal subsets. Average gene expression is shown by color intensity. Percent of cells expressing each gene is shown by size. **C&D.** Differential abundance beeswarm plots from LP1/2 (**C**) and MP (**D**) layers by cell type. Each dot is a neighborhood of cells calculated using miloR. Neighborhoods that reach significance (spatial FDR < 0.1) are colored by log fold-change. **E**. Bar plot representing the total number of up-(red) or down-regulated(blue) genes by cell type in the LP1/2 layers **F.** Bar plot representing the total number of up-(red) or down-regulated(blue) genes by cell type in the MP layer.

Given the above results, we next interrogated transcriptional changes within the stromal niche. Compatible with the global transcriptional view in Figure 1G&H, the LP1/2 layer contained the stronger perturbation of gene expression within each stromal cell type compared to the MP layer. Both CXCL14+ fibroblast populations as well as adventitial PI16+ fibroblasts had a large number of significantly differentially expressed genes when comparing CDs to CDni and NL, suggesting their contribution to stricture (Figure 2E). The stromal transcriptional changes in the MP layer were minimal, but most pronounced in CDs compared to NL segments. Cell types contributing to transcriptional changes were mainly stem-like fibroblasts and ACTA2+ smooth muscle cells (Figure 2F). The per cell type signaling pathways can be found in Supplementary Results and Supplementary Figure 3A&B. We further mapped previously published stromal populations associated with IBD onto our dataset and interrogated the frequency of the cell types per segment. These results can be found in Supplementary Results and Supplementary Figure 4A&B.

To understand global communications and interactions among cells we used CellChat^14^ to quantitatively infer intercellular communication networks. CellChat predicts major signaling inputs (incoming receptor signals, received signals) and outputs (sent signals, outgoing signals) on a per cell type basis. Our initial analysis pointed to CDi and CDs as the most relevant areas underlying the progression from non-strictured inflammation to stricture. In LP1/2 the major populations with strong incoming receptor signals in CDs compared to CDi are myeloid cell types (CLCL9/CXCL10^high^, inflammatory monocytes, NLRP3^high^ and Metallothionein^high^), T/NK cells (CCL5/CD8+, CCL4/CCL5/CD8+, XCL1+, CCL5+), endothelial venular cells, MMP/WNT5A+ fibroblasts and pericytes (Figure 3A). In MP fraction, major populations that show strong incoming receptor signals are B cells (CCL3/4+ plasma cells, mature B3/CRIP1+, CYBA/IGHG4+ plasma cells), myeloid cells (cDC2, MEG3/MEG8^high^, Metallothionein^high^) and stromal cells (MMP/WNT5A+) (Figure 3C). Strikingly, fibroblasts stand out as major signal senders in CDs compared to CDi in LP1/2 layer, particularly CXCL14/ADAMDEC1+, MMP/WNT5A+, CXCL14/F3/PDGFRA+ and ECM^high^ cells (Figure 3B). A comparable finding was observed in the MP layer, with pericytes and SMC/HHIP+ stromal cells showing stronger outgoing signals in CDs compared to CDi. All other cell fractions had a negligible contribution as signal senders (Figure 3D). When evaluating the outgoing signal strength across segments in the stromal compartment the noted changes were found to be progressive with fibroblasts taking on an increasing signal sending role from NL, over CDni, CDi to CDs (Figure 3E, Supplemental Figure 5). These findings indicate that fibroblasts are major ‘signal sending hubs’ in CDs in the LP1/2 as well as MP layers.

**Figure 3.**
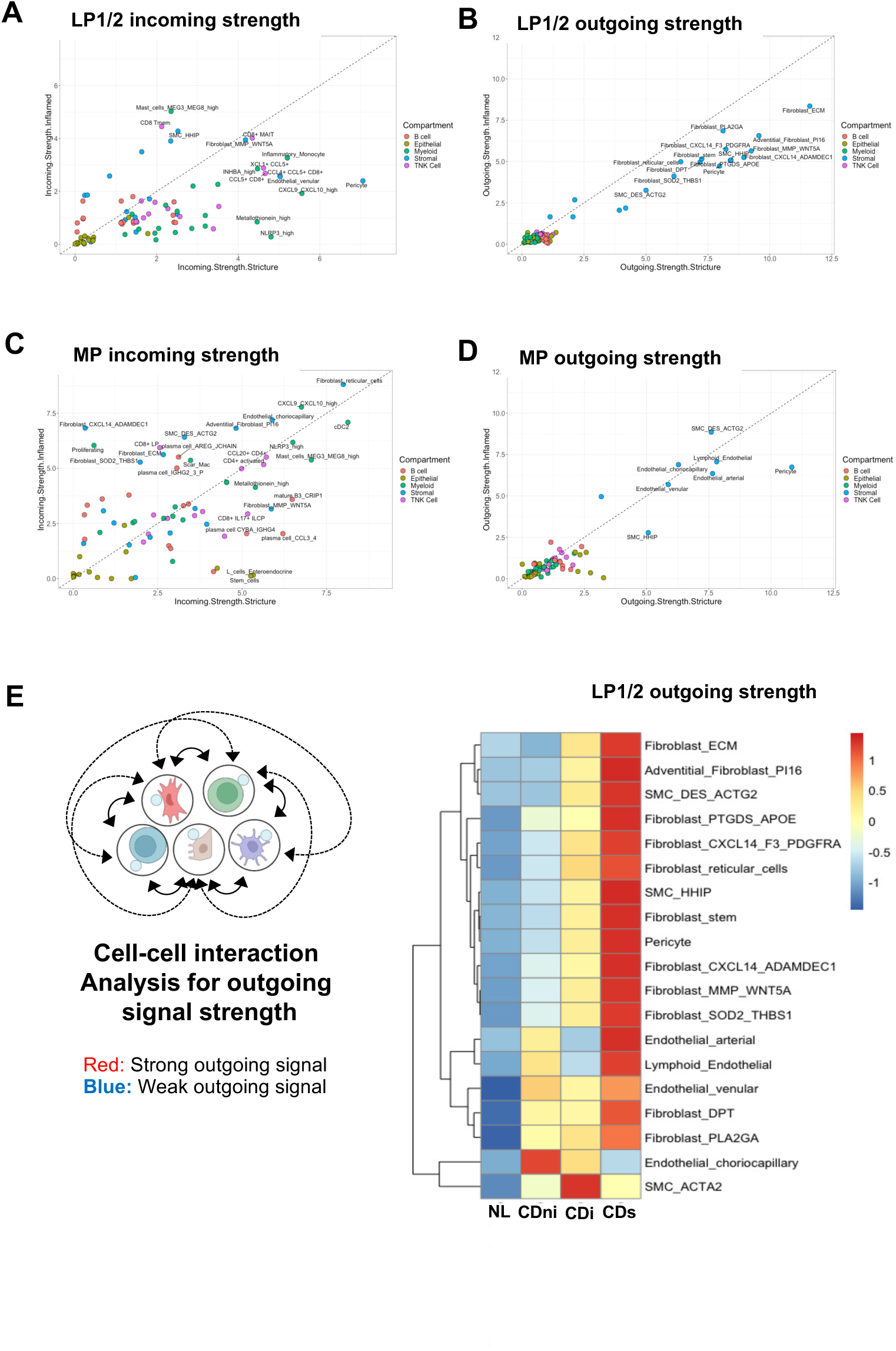
Cell-cell interaction analysis of the full scRNAseq atlas by tissue segment and layer. **A&B.** Scatterplot of the incoming (**A**) and outgoing (**B**) interaction strength for CDs versus CDi sections in the LP1/2 layers calculated using cellChat^14^. Each dot represents a cell type **C&D.** Scatterplot of the incoming (**C**) and outgoing (**D**) interaction strength for CDs versus CDi sections in the MP layers calculated using cellChat. Each dot represents a cell type. **E**. Heatmap of outgoing signal strength of the stromal compartment by cell type and segment, scaled by cell type. Red indicates high and blue low signal strength.

### Cadherin 11 emerges as a dominant cell surface molecule in Crohn’s disease strictures

Our cell-cell interaction analysis revealed a dominant role for fibroblasts as pathobiologically relevant ‘signal senders’ as well as ‘receivers’. We hence queried molecules that may mediate fibroblast-fibroblast interactions by focusing on three populations that were overrepresented in the LP1/2 compared to MP layers and either increased in CDs segments (MMP/WNT5A+) or showing major transcriptional changes in CDs (CXCL14/PDGFRA+ and CXCL14+/ADAMDEC1+). When interrogating overlapping gene signatures of these populations, we detected 12 shared markers (Figure 4A&B), of which only *CDH11* is expressed on the cell surface and in addition is highly selective to fibroblasts (Supplemental Figure 4C). When analyzing *CDH11* expression across stromal cell types, LP1/2 layer neighborhoods that had a high *CDH11* expression also had an increased abundance or differentially expressed genes in CDs, making this a marker for stromal cells enriched in CDs (Figure 4C). A shift in the density of *CDH11* expression is noted in parallel to disease progression from NL, CDni over CDi to CDs (Figure 4D). Of relevance, CDH11+ stromal cells are not only increased in abundance in CDs, but *CDH11* is increased in CXCL14/ADAMDEC1+ fibroblasts on a per cell basis (Figure 4E). Automated cyclic immunofluorescence analysis on FFPE intestinal tissue sections confirmed robust upregulation of CDH11 in the mucosa/submucosa in CDs with minor expression in the MP. CDH11 was co-expressed with αSMA and desmin (DES) and the frequency of CDH11/αSMA/DES+ cells compared to total cells or within the αSMA/DES+ fraction increased in CDs (Figure 4F&G).

**Figure 4.**
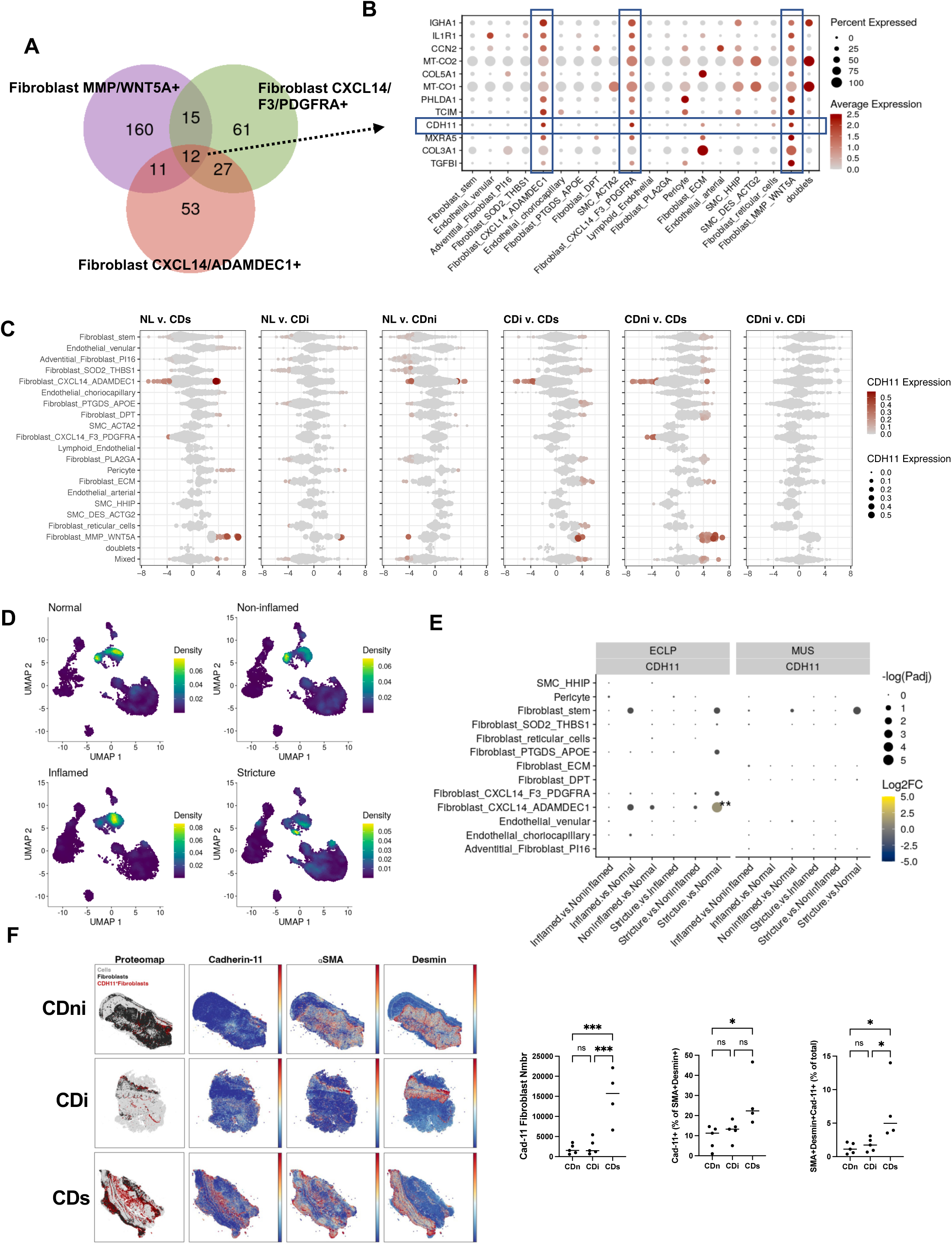
*CDH11* is a potential regulator of intestinal fibrosis. **A**. Venn diagram showing overlap of significant markers for CXCL14/ADAMDEC1+, CXCL14/F3/PDGFRA+, and MMP/WNT5A+ fibroblasts. **B.** Dot plot of 12 overlapping markers from A. **C.** Differential abundance beeswarm plots from miloR analysis of LP1/2 layers in stromal compartment with neighborhoods colored by *CDH11* expression. Only significantly differentially abundant neighborhoods show *CDH11* expression. **D**. UMAP plots of *CDH11* expression density in stromal compartment split by segment (NL, CDni, CDi, and CDs). **E**. Dot plot demonstrating *CDH11* pseudobulk differential expression by cell type. Significance is shown by size and denoted by ** (padj :< 0.01). Only contrasts with a significant adjusted p-value (padj :< 0.05) show log fold change color. **F**. Cyclic immunofluorescence of intestinal FFPE section using vimentin, aSMA and DES antibodies. Automated quantification is indicated. Data are presented as mean ± SD. *p<0.05; p<0.001.

### Cadsherin-11 is upregulated in IBD tissues

Underscoring the pathophysiological relevance of CDH11 in CDs, we found upregulation of its gene and protein expression in full thickness CDs, CDns and UC compared to the NL tissues (Figure 5A&B), with the most pronounced increase (up to 35-fold) found in CDs. Dual-immunolabeling with markers of mesenchymal, endothelial and epithelial cells and leukocytes co-localized CDH11 with α-SMA and vimentin in the mucosa and submucosa, and enrichment in a sub-fraction of desmin-positive cells (Figure 5C and Supplemental Figure 6) compatible with high CDH11 expression by myofibroblasts. Expression of CDH11 was low in the muscularis propria and did not change based on the disease phenotype (data not shown). Minimal to no co-localization of CDH11 was found with CD45, CD68, E-cadherin and CD31 (Supplemental Figure 6), indicating that immune cells, including tissue resident macrophages, are minimal contributors to CDH11 expression in the inflamed gut.

**Figure 5.**
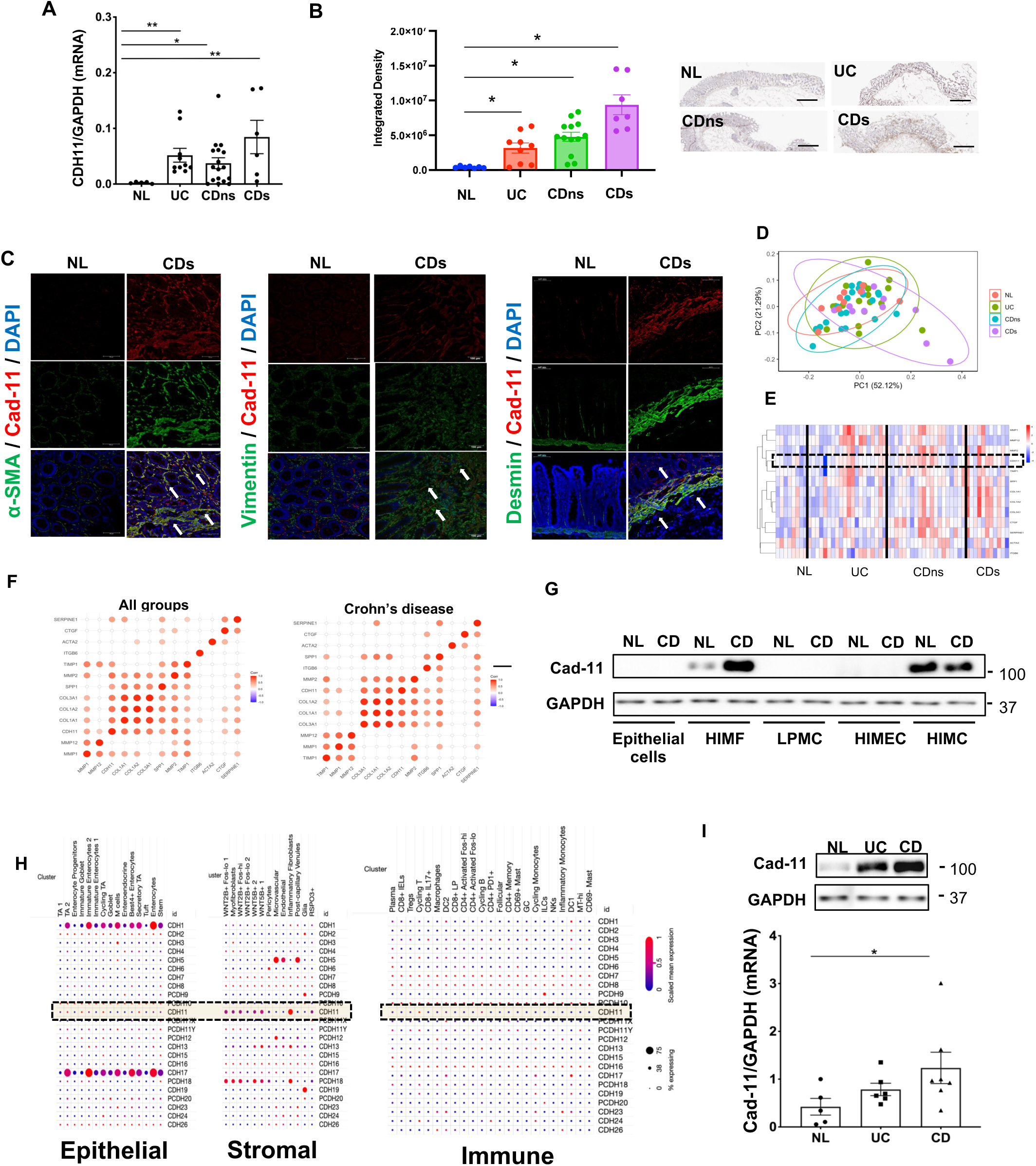
Cadherin-11 expression is upregulated in the intestine of IBD patients. **A**. Expression of *CDH11* mRNA in fresh full thickness sections. *CDH11* level was normalized by GAPDH mRNA expression (n = 43). **B.** Immunohistochemistry of full thickness intestinal resection tissues immunolabelled with Cadherin-11 (CDH11) and quantified by integrated density values (IDV). **C.** Full thickness sections of NL and CDs patients were dual-immunolabeled for CDH11 (red) and either α-SMA, vimentin, or desmin (green). Representative confocal images with the full panel being available in Supplemental Figure 6. Arrows point to colocalization of CDH11 with myofibroblast markers. Representative for n=4 per group. **D&E.** Principal components analysis and heatmap analysis of the relative abundance of ECM related gene expression in FFPE tissue using a pre-specified NanoString nCounter panel and normalized to housekeeping genes. Ellipses indicate 95% confidence intervals. Separation of the gene panel was noted in CDs compared to the other groups (n = 59). **F.** Correlograms showing statistically significant correlations for relative abundance of ECM related gene expression in FFPE tissue using a pre-specified NanoString nCounter panel (Spearman correlation, P ≤ .05). Red dots indicate positive correlation. **G.** Expression of CDH11 was examined by immunoblotting analysis in the following primary human intestinal cell types: Intestinal epithelial cells; HIMF, human intestinal fibroblasts; LPMC, lamina propria mononuclear cells; HIMEC, human intestinal microvascular endothelial cells; HIMC, human intestinal muscle cells. **H.** Gene expression of known cadherins in the epithelial, stromal and immune compartment, using a publicly available single cell RNA sequencing dataset from UC patients.^7^ **I.** Expression of CDH11 protein in HIMF isolated from CD, UC patients and non-IBD control was compared by immunoblotting analysis and densitometric quantification, Data are presented as mean ± SD (n=17). Data are presented as mean ± SD (n = 8-12). *: *p*<0.05.** *p*<0.01. Abbreviations: NL, normal; UC, ulcerative colitis; CDns, Crohn’s disease, non-strictured; CDs, CD strictured.

A pre-specified NanoString panel comprising 13 genes associated with fibrotic diseases in an independent cohort of formalin-fixed paraffin-embedded (FFPE) full thickness intestinal tissues revealed overlap between NL, UC and CDns in a principal component analysis and heatmap. Some genes expressed in CDs were separated from the other groups, compatible with upregulation of *CDH11* (Figure 5D&E). Across all IBD and control samples and within CD samples, *CDH11* was positively correlated with COL1A1, CvOL1A2, COL3A1, SPP1, TIMP1 and SERPINE (Figure 5F).

### IBD myofibroblasts express increased levels of cadherin-11

Investigation of various human intestinal cells revealed that CDH11 was expressed by HIMF and human intestinal muscle cells (HIMC), but not epithelial cells, lamina propria mononuclear cells (LPMC) or human intestinal microvascular endothelial cells (HIMEC) (Figure 5G). Analysis of a publicly available scRNAseq dataset^7^ from UC mucosal biopsies confirmed that CDH11 mean and percent expression clusters in the stromal compartment, with the strongest signal in inflammatory fibroblasts. The epithelial and immune cell compartments did not express CDH11 (Figure 5H). CDH11 expression was higher in CD HIMF compared to NL (Figure 5I).

### Modulation of cadherin-11 activity affects myofibroblast ECM production

We next investigated the effect of CDH11 loss-of-function in HIMF with blocking antibody H1M1^15^ or siRNA knockdown (KD). We developed a novel medium throughput ECM deposition assay where the deposited (versus soluble or gene expression) ECM is automatically measured (*Methods)*. Treatment with H1M1 reduced deposition of FN and ColI/III in a dose-dependent manner in NL, UC, CDns and CDs HIMF (Figure 6A&B). CDH11 KD markedly downregulated its expression and decreased cellular level of FN (Figure 6C). In contrast, CDH11 gain-of-function by extracellular recombinant human CDH11 domain-Fc chimera (hCDH11-Fc) (‘CDH11 activator’)^16^ resulted in dose-dependent increase of FN and ColI/III deposition by NL, UC, CDns and CDs HIMF (Figure 6A&B), an effect comparable to TGF-β1, the strongest profibrotic activator of HIMF (Figure 6B). CDH11 activator only affected HIMF function when it was plate-bound but not in soluble form (data not shown). Given the analysis of scRNAseq suggesting a dominant signal sender role of CDs fibroblasts, we analyzed the HIMF secretome with and without CDH11 activation (Figure 6D&E, Supplemental Figure 7, Supplemental Table 7). 332 proteins were significantly different upon CDH11 activation, with 179 up- and 153 down-regulated. CDH11 activation upregulated secretion of multiple core matrisome proteins suggesting a broad pro-fibrotic effect. Pathway enrichment analysis indicated the upregulation of multiple proteins and signaling pathways related to inflammation, suggesting a pro-inflammatory effect of CDH11 (Figure 6E). The detailed analysis of the secretome is found in Supplemental Figure 7 and Supplemental Table 7.

**Figure 6.**
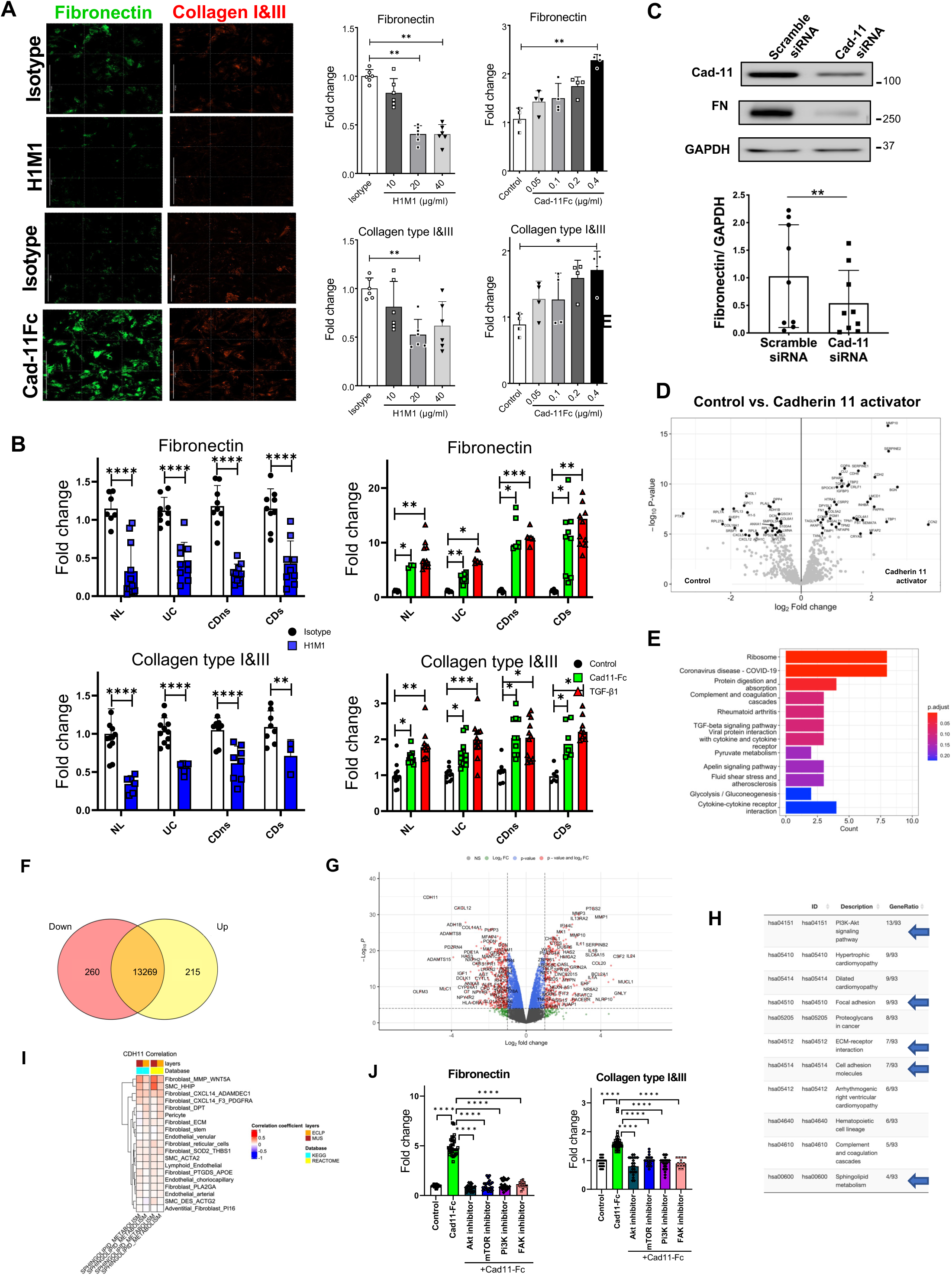
Cadherin-11 regulates the fibrogenic phenotype of human intestinal myofibroblasts. CDH11 was either inhibited in HIMF isolated from subjects using CDH11 blocking antibody (H1M1) or specific siRNA to CDH11 or activated by CDH11 activator (Cad11-Fc). **A.** HIMF from normal (NL) intestinal tissue were exposed to different doses of H1M1 or Cad11-Fc and extracellular matrix deposition (FN and ColI/III) was assayed using the ECM deposition assay (n=6 per group). Representative immunofluorescence images are shown. Scalebar=1000 µm. **B.** HIMF from the patient groups NL, normal; UC, ulcerative colitis; CDns, Crohn’s disease, non-strictured; CDs, CD strictured were exposed to H1M1 (20µg/ml) or Cad11-Fc (0.4µg/ml), respectively and assayed using the ECM deposition assay. TGF-β1 (1 ng/ml) was used as a positive control (n=12). **C.** Knockdown of CDH11 was performed in HIMF and CDH11 and FN expression was determined by immunoblotting and densitometric quantification (n=18). **D.** Annotated volcano plot of secreted proteins with upregulated (right) and downregulated (left) proteins with and without CDH11 activation in HIMF NL (n=9 for each group). **E.** Protein enrichment analysis of the HIMF NL secretome in response to CDH11 activation. **F.** Significantly up (yellow) and downregulated (red) genes upon *CDH11* siRNA knockdown (KD) in HIMF NL (n=4 each for the scrambled control and knockdown groups). **G.** Annotated volcano plot with upregulated (right, red) and downregulated (left, red) genes upon *CDH11* KD. **H.** KEGG pathway enrichment analysis upon *CDH11* KD. **I.** Heatmap of the correlation of CDH11 normalized expression with sphingosine signaling. Ucell scores derived from KEGG and REACTOME databases in mesenchymal cell types in our scRNAseq dataset. **J.** HIMF NL were exposed to hIgG1 (control) or Cad11-Fc (0.4µg/ml) with or without compound inhibitors (GSK690693 (Akt inhibitor, 5µM); mTOR Inhibitor XI (mTOR Inhibitor, 0.25µM); LY-294,002 hydrochloride (Pi3K inhibitor, 30µM); Y15, 1,2,4,5-benzenetetramine tetrahydrochloride (FAK inhibitor, 5µM) and assayed using the ECM deposition assay (n=18). Data are presented as mean ± SD. *p<0.05;**p<0.01; ***p<0.001; ****p<0.0001.

### Global gene expression mediated by cadherin-11

Next generation RNA sequencing (RNAseq) in NL HIMF with and without gene KD of *CDH11* resulted in differential expression of 475 genes (260 downregulated, 215 upregulated) (Figure 6F&G). The top up- and down-regulated genes, correlation analysis and gene enrichment analysis with *CDH11* confirm correlation of *CDH11* with ECM deposition, remodeling, and cell proliferation (Supplemental Figure 8A to C, Supplemental Tables 8&9). *CDH11* strongly correlated with a majority (116/161) of known matrisome genes including those encoding fibronectin (FN1), tenascins (TNXB/C), ECM1/2, latent transforming growth factor beta binding proteins (LTBP1/2/3), transforming growth factor beta-induced (TGFBI), and several collagens (COL1, COL3-6). *CDH11* knockdown reduced expression of multiple proinflammatory genes (IL1B, TNF). Among modulation of multiple pathways, *CDH11* KD revealed a robust downregulation of sphingosine and FAK signaling (Figure 6H, Supplemental Tables 8 to 10). As a result of this data and to validate scRNAseq as a tool for therapeutic target discovery, we chose to interrogate our stromal populations of interest to CDs (CXCL14+ and WNT5A+ fibroblasts) in our scRNAseq dataset. Corroborating the NGS results, sphingosine metabolism was strongly linked with *CDH11* expression in these populations (Figure 6I). Modulation of sphingosine and FAK signaling pathways by inhibition of AKT, mTOR, Pi3K and FAK with selective small molecules, led to amelioration of CDH11-induced ECM deposition, functionally confirming the relevance of these pathways to CDH11 pro-fibrotic function (Figure 6J).

### Functional ablation of cadherin-11 improves experimental intestinal fibrosis

We lastly investigated the therapeutic potential of CDH11 inhibition *in vivo* by using a total knockout of Cdh11 (Cdh11-KO) in the DSS-induced fibrosis model of colitis.^15^ The clinical score was reduced in Cdh11-KO throughout the course of DSS administration (Figure 7A). Cdh11-KO showed no difference in inflammation compared to WT mice (Figure 7B), but reduced picrosirius red, ColI, FN staining (Figure 7C) and reduced thickening of the muscularis mucosa, submucosa and muscularis propria (Figure 7D). Cdh11-KO mice displayed reduction in gene expression of *IL6* and *TNF* compared to WT mice (Figure 7E).

**Figure 7.**
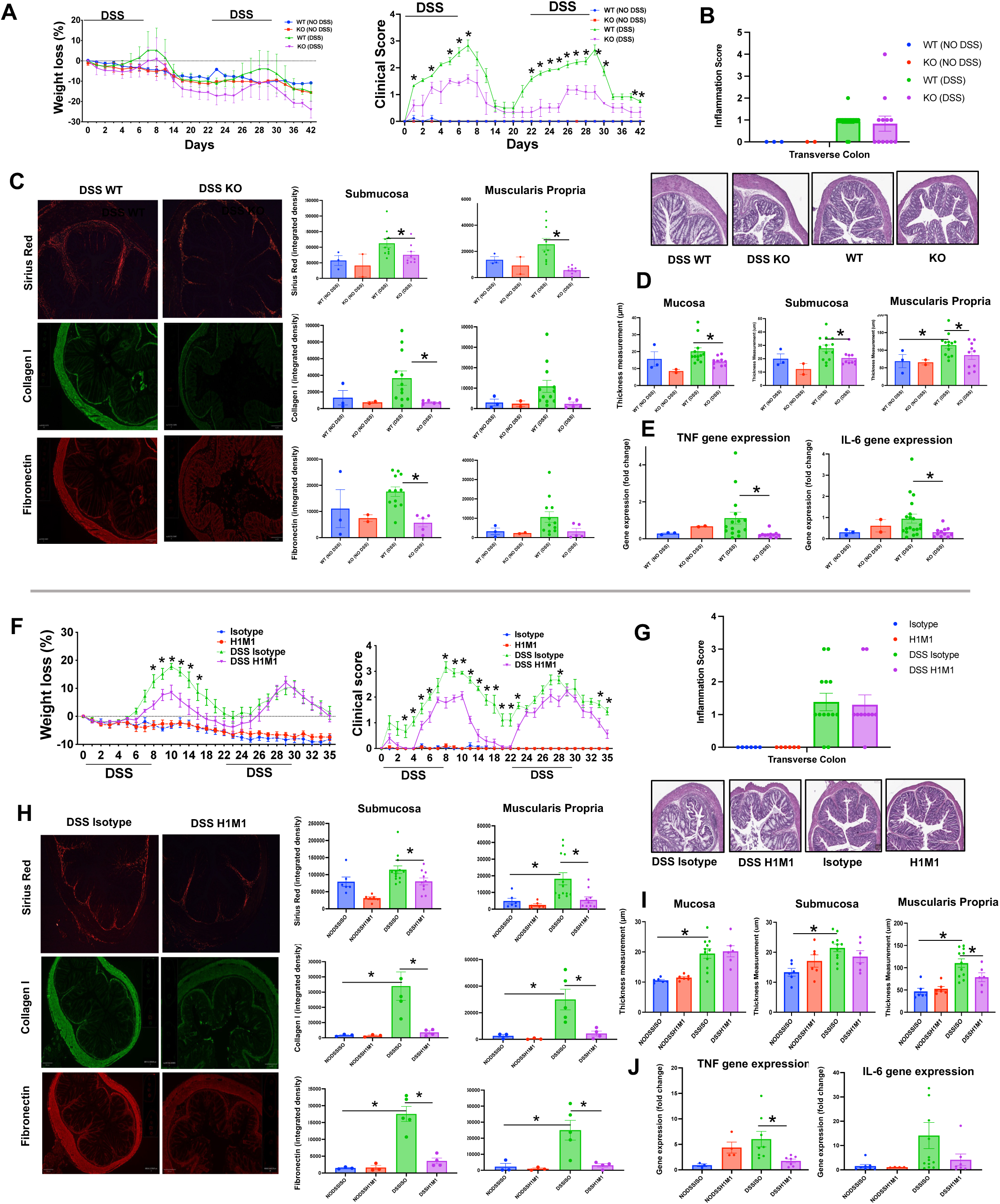
Blockade or genetic ablation of cadherin-11 attenuates chronic DSS induced colitis in mice. Cdh11 null (KO) mice and wildtype (WT) littermates were subjected to two cycles of dextran-sodium sulfate (DSS) administration (2%) followed by post-DSS recovery. **A.** The severity of DSS induced colitis was evaluated by measuring the body weight loss and calculating the clinical score (n=12 per group). Colonic sections were fixed and stained with hematoxylin and eosin (H&E) for histological examination. **B**. The inflammation score was determined using H&E sections by an IBD pathologist in a blinded fashion. **C.** The severity of fibrosis was evaluated through Sirius red staining, Col1 or FN immunolabeling and quantified using integrated density measurements for the submucosa and the muscularis propria separately. **D.** Thickness of the intestinal layers was measured for mucosa, submucosa and muscularis propria. **E.** Gene expression of TNF and IL6 in intestinal tissues. Wildtype (WT) mice were subjected to two cycles of dextran-sodium sulfate (DSS) administration (2%) followed by post-DSS recovery. Cdh11 blocking antibody H1M1 or isotype antibody were given daily starting day 1 of the experiment. **F.** Severity of DSS induced colitis was evaluated by measuring the body weight loss and calculating the clinical score (n=10 per group). **G**. Inflammation score was determined using H&E sections by an IBD pathologist in a blinded fashion. **H.** Severity of fibrosis was evaluated through picrosirius red staining, Col1 or FN immunolabeling and quantified using integrated density measurements for the submucosa and the muscularis propria separately. **I.** Thickness of the intestinal layers was measured for mucosa, submucosa and muscularis propria. **J.** Gene expression of TNF and IL6 in intestinal tissues. Data are presented as mean ± SEM \**p*<0.05.

We then examined the effect of CDH11 blocking antibody H1M1 in DSS fibrosis. H1M1 pharmacokinetics (PK) study established the dose of 0.5 mg/mouse every other day as optimal (Supplemental Figure 9), reaching a minimum plasma H1M1 level >200μg/ml to achieve optimal efficacy^17^. Upon preventive administration of H1M1 the clinical score was reduced during DSS administration and post-DSS recovery (Figure 7F). H1M1 did not reduce severity of inflammation^18^ (Figure 7G), but reduced development of fibrosis as measured by picrosirius red, FN1 and ColI staining (Figure 7H) and reduced muscularis propria thickening (Figure 7I). H1M1 also reduced gene expression of *TNF* and showed a trend for reduced *IL6* expression (Figure 7J).

## DISCUSSION

Organ fibrosis is a universal response to multiple pathological conditions of inflammatory, neoplastic or degenerative nature.^5^ Depending on its extent and location, fibrosis has serious consequences that become the primary cause of symptoms and disease evolution as well as choice of therapeutic intervention. This is well exemplified by the management of fibrosis-induced stricture formation in CD, which is still very challenging due to a limited understanding of the underlying cellular and molecular mechanisms.^2^ To overcome this limitation, we analyzed the transcriptome of 409,001 intestinal cells from 20 patients and generated the first full thickness CDs scRNAseq atlas. Our dataset provides novel information on the epithelial, immune and stromal compartments and enables the characterization of cell type specific differences along tissue segments and layers. These data point to *CDH11* as a key driver of stricturing CD and a putative therapeutic target.

Our results contribute several new key findings for CDs. First, the major differences between non-inflamed, inflamed and strictured bowel segments at the cell type and transcriptional level are found in the LP1/2 fraction, reflecting the mucosa and submucosa. Far fewer alterations were detected in the MP of CDs despite its marked thickening and contribution to obstructive symptoms.^2^ This may be explained by post-transcriptional regulation of HIMC proliferation and activation or by the use of tissue samples in a late process of stricture formation. Various stromal cell types were exclusive to the LP1/2 or MP fractions, which highlights the unique features of each tissue layer.

Second, we identified fibroblasts as major signal sending hubs in CDs. This is highly relevant as to date it was unclear which mechanisms drive the progression from pure inflammation to fibrostenosis and whether progression is driven by inflammation-dependent or -independent processes.^2^ Being major signal senders, this puts fibroblasts in the center of strategies to block transition of inflammation to fibrosis or directly preventing or eliminating fibrosis.

Third, our work identified in CDs an abundance of MMP/WNT5A+ fibroblasts, a cell type expressed exclusively in the LP1/2 but not the MP fractions. This fibroblast type matches a cell known as inflammatory fibroblast (IL13RA2/IL11+)^7^ from the mucosa of patients with UC resistant to anti-TNF treatment. One could speculate that the presence of MMP/WNT5A+ and the reported association with anti-cvTNF resistance is mediated by the presence of fibrosis. We also found a CXCL14+ fibroblast population co-expressing either PDGFRA or ADAMDEC1 displaying the most pronounced transcriptional changes in CDs, and hence of functional relevance. CXCL14+ fibroblasts have been associated with increased ECM production^19^ and CXCL14 itself has profibrotic properties.^20^ ADAMDEC1 is critical in the fibroblastic response to tissue remodeling and healing during colitis^21^ and PDGFRA+ fibroblasts display a proliferative phenotype and are a major source of collagen.^22, 23^

Based on the prominent role of fibroblasts in CDs signaling, the differences largely restricted to the LP1/2 fraction, the over-representation of distinct fibroblast types, and the strong contribution to transcriptional changes in CDs, we sought a unifying molecule and identified CDH11 as the only cell surface receptor shared by the CXCL14+ and MMP/WNT5A+ fibroblasts. While also expressed in macrophages, CDH11 is primarily responsible for mediating homophilic stromal cell-cell interactions,^24–27^ is upregulated in fibrotic disorders of the lung,^24^ liver,^25^ skin,^26^ intestine,^27^ and its inhibition attenuates fibrosis in multiple animal models.^25^ This raises the prospect of using CDH11 blockade to prevent or treat CD-associated intestinal fibrosis.

In support of the above, we confirmed in CDs a marked upregulation of CDH11 mRNA and protein expression,^27^ and correlation of CDH11 gene expression with hallmark profibrotic genes. CDH11 was predominantly expressed in fibroblasts in bowel tissue, isolated primary intestinal cells, and in scRNAseq. CDH11+ cells were more frequent in CDs and on a per cell basis CDH11 was upregulated in MMP/WNT5A+ inflammatory fibroblasts. By using a novel ECM deposition assay we also showed that CDH11 has strong pro-fibrotic properties *in vitro* comparable in magnitude to those of TGF-β1. Measuring actual ECM deposition instead of gene expression is important as it most closely reflects the *in vivo* situation. Using proteomics and next generation sequencing we uncovered not only a broad pro-fibrotic effect of CDH11 *in vitro*, but also its additional previously underappreciated pro-inflammatory property and the potential mechanism of sphingosine and FAK signaling, a finding which was confirmed in our scRNAseq dataset and functionally validated using small molecule inhibitors in HIMF.

Importantly, a pharmacological (antibody-mediated) or genetic inhibition of CDH11 decreased intestinal fibrosis *in vivo* without any adverse effects after prolonged administration. This is consistent with previously published data in TNBS colitis in CDH11-KO mice^27^ and in addition to providing a different fibrosis model adds the successful administration of a blocking antibody. In our experimental fibrosis models however, we were only able to appreciate the anti-inflammatory properties on the intestinal gene expression level for *TNF* and *IL6*, but not on histopathologic inflammation scoring. This may indicate that the anti-fibrotic effects outweigh the anti-inflammatory effects *in vivo*.

In conclusion, by developing a detailed single cell atlas of non-inflamed, inflamed and strictured autologous CD tissue segments, characterizing the strong pro-fibrotic activity of specific mesenchymal cell subsets in the mucosa and submucosa, and identifying CDH11 as a widely expressed cell surface molecule essential for ECM production, we uncovered novel cellular and molecular mechanisms involved in CDs and CDH11 as a potential therapeutic target.

## Supporting information

Supplemental Details and Figures

## Grant support

This work was supported by the Helmsley Charitable Trust through the Stenosis Therapy and Anti-Fibrotic Research (STAR) Consortium (to F.R.), the Crohn’s and Colitis Foundation Fellowship Award (No. 648334 to J.L.), the National Institute of Health (R01DK123233 to F.R., P30 DK097948 to C.F. & F.R.) and Pliant and Pfizer Inc. through sponsored research agreements. No sponsor had any involvement in study design, collection or interpretation of data. Pfizer Inc. supported bioinformatic analysis in collaboration with the Cleveland Clinic.

## Conflict of Interest

P.K.M., Q.T.N., J.L., S.Z., G.A.W., J.C., S.L., J.W., R.M., P.K., T.P., S.H., N.N., M.D., S.T., S.R.S,

A.I.I have no conflict of interest.

I.O.G. received funding from UCB, Celgene, Gossamer and Pliant.

S.D.H. was consultant to Shionogi and Takeda.

S.M.C., S.L., T.F., S.A., K.B., K.H., R.M., K.D., T.W., and K.M.K. are employees of Pfizer, Inc and may hold stock equity.

C.P. was a visiting graduate scholar at Pfizer, Inc.

P.K., M.D., S.T. are employees of Pliant Inc.

C.F. received speaker fees from UCB, Genentech, Sandoz, Janssen and he is consultant for Athos Therapeutics, Inc.

F.R. was consultant to AbbVie, Adnovate, Agomab, Allergan, Arena, Astra-Zeneca, Boehringer-Ingelheim, Celgene/BMS, CDISC, Celsius, Cowen, Ferring, Galapagos, Galmed, Genentech, Gilead, Gossamer, Granite, Guidepoint, Helmsley, Horizon Therapeutics, Image Analysis Limited, Index Pharma, Landos, Jannsen, Koutif, Mestag, Metacrine, Mopac, Morphic, Organovo, Origo, Pfizer, Pliant, Prometheus Biosciences, Receptos, RedX, Roche, Samsung, Sanofi, Surmodics, Surrozen, Takeda, Techlab, Teva, Theravance, Thetis, UCB, Ysios, 89Bio.

## ABBREVIATIONS

CD: Crohn’s disease
CDi: non-strictured but inflamed CD
CDni: non-strictured, non-inflamed CD
CDs: strictured CD
CDH11: Cadherin-11
DMEM: Dulbecco’s modified Eagle medium
DSS: Dextran sodium-sulfate
ECM: Extracellular matrix
FAK: focal adhesion kinase (FAK)
FBS: Fetal bovine serum
FFPE: Formalin-fixed paraffin embedded
HIMFs: Human intestinal myofibroblasts
IBD: Inflammatory bowel diseases
KO: knockout
LP: lamina propria
MP: muscularis propria
NL: Non-IBD patients
PBS: Phosphate-buffered saline
scRNAseq: single cell RNA sequencing
SES-CD: Simple Endoscopic Score for Crohn’s Disease
UC: Ulcerative colitis
UMI: Unique Molecular Identifier
WT: wild type

## AUTHOR CONTRIBUTION

Study design: PKM, JL, SC, GAW, FR; Execution/data collection: PKM, QTN, JL, SZ, SC, GAW, JC, IOG, SL, JW, RM, PK, TP, SL, DC, CR, TF, SA, KB, KH, CP, KD, SH, NN, MD, SDH, SRS;

Data compilation and analysis: PKM, QTN, CP, SA, SL, SC, JL, FR; Oversight/advisory: PKM, SC, TW, ST, RM, KMK, FR; Wrote and edited manuscript: all authors; Acquired funding, regulatory approvals: JL, CR, FC, FR

## ACKNOWLEDGEMENTS

We thank Dr. Judith Drazba, Dr. John Peterson, Andrelie Branicky, Apryl Helmick and Bridget Besse from the LRI Microscopy and Image Core of the Cleveland Clinic for help in confocal microscopy and image analysis. We thank Dr. Belinda Willard and Dr. Isaac Ampong from the Metabolomic and Proteomics Core of the Cleveland Clinic for support of the proteomics analysis. We appreciate the support of the Genomics Core of the University of Chicago for next generation sequencing. We thank Pfizer Inflammation and Immunology Research Units’s Fibrosis Discovery Group and Systems Immunology group for helpful comments and support, as well as Pfizer’s cellular and optical microscopy technology center for their imaging support. We acknowledge the support of the Departments of Colorectal Surgery and Pathology of the Cleveland Clinic. Tissue samples were provided by the Human Tissue Procurement Facility of the Cleveland Clinic through the services of the Biorepository Core funded by P30 DK097948-06. We thank the contribution of the Cleveland Clinic Center for Global Translational IBD. We also deeply appreciate the contribution of patients who donated the tissues.

