## Supplemental Details and Figures for "Stricturing Crohn’s disease single-cell RNA sequencing reveals fibroblast heterogeneity and intercellular interactions"

#### SUPPLEMENTAL MATERIALS

#### SUPPLEMENTAL METHODS

*Tissue dissociation and isolation of single cells*

A two-step collagenase/dispase digestion-based method was used to isolate single cells from the epithelium, lamina propria (LP), submucosa and muscularis propria (MP) from all tissue groups. Each sample was mechanically dissected to separate the epithelium, LP and submucosal layers from the MP layer as previously described.[1, 2] The epithelium, LP and submucosal layers were incubated in 0.15% dithiothreitol (DTT, Sigma, St. Louis, MO) and stirred for 30 minutes, followed by digestion with an enzyme mix containing collagenase-I (3 mg/mL, Worthington, Lakewood, NJ) and dispase (3 mg/mL, Roche, Basel, Switzerland), for 30 min at 37 °C without stirring. The resulting cell suspension containing a mixture of epithelial cells and LP mononuclear cells was filtered through a 100 µm cell strainer to retain the undigested tissue fragments and collect the epithelial and LP cell enriched fraction (termed "LP1" fraction) The retained tissue was subjected to a second digestion with a fresh same enzyme mix for another 30 minutes, and then filtered through a 100 µm cell strainer to obtain a "LP2" fraction. MP segments from the same tissue were mechanically minced and subjected to digestion with collagenase-II (2 mg/mL, Worthington, Lakewood, NJ) for 60 minutes at 37 °C, mixing every 15 minutes by inversion. The tissue digest was filtered through 100 µm filter. The resulting cell suspension was spun and collected, while the remaining tissue digest was incubated again for 30 min in enzyme-free complete medium at 37 °C and filtered through 100 µm strainer to obtain a second cell suspension. The two MP cell suspensions were combined and used as a single MP cell fraction. The isolated

LP1, LP2 and MP cell fractions were assessed for viability with Trypan blue, counted, aliquoted, fixed in cold 80% methanol and stored at –80 °C until use. Quality metrics for single cell isolation were presence of single cells with no or minimal clumps and viability greater than 80%.

*Library generation*

Methanol-fixed cells from mucosa/sub-mucosa single cell suspensions were thawed, rehydrated, and immediately processed using either the 3' v3.1 or 5' v2 chemistries as described below on the 10X Chromium single cell platform (10X Genomics). Chemistry type for each sample is included in Supplemental Table 1. For 3' v3. 1 samples, GEMs were generated from cells with reverse transcription master mix, gel beads, and oil loaded on a Chromium Next GEM Chip G. For 5' v2 samples, GEMs were generated from cells with reverse transcription master mix, gel beads, and oil loaded on a Chromium Next GEM Chip K. After cDNA amplification, libraries were prepared and single-indexed with Single Index Plate T Set A. Libraries were sequenced on either a NextSeq or a NovaSeq, according to manufacturer's instructions.

*Single-cell RNA analysis*

After sequencing, samples were demultiplexed using bcl2fastq and fastq files were aligned to the human genome (GRCh38). Sample analysis was performed using the R package Seurat. Ribosomal protein genes were removed and cells were filtered at 200 genes per cell and mitochondrial percentage < 30. Counts were normalized using SCTransform (3' and 5' normalized separately) and mitochondrial percentage and nCount_RNA were regressed out. Harmony was used to correct for patient and chemistry variation. For the global compartment the data was split by chemistry and normalized separately. The use of the first 50 PCs and a resolution of 0.1 was used to identify broad cell types in the global compartment. For mesenchymal subclustering, the data was split by chemistry and normalized separately. Highly variable genes from each chemistry were merged (4070 total genes) and use of the first 25 PCs and a resolution of 0.4 revealed multiple non-mesenchymal clusters. These were removed and the process was repeated with 4072 genes, 30 PCs and a resolution of 0.6. Cell annotation was performed manually using marker genes and known canonical markers. Pseudobulk differential expression for each cell type or subtype was performed using DESeq2, with samples containing a minimum of 20 cells in the cell type and genes expressed in at least the minimum contrast group size. Differential abundance was performed using miloR with k = 30, d = 25, a cell type fraction of 0.7, and spatial FDR of 0.1. In order to characterize the cell-cell interactions, each of the compartments were merged and doublet populations were removed. We divided the merged compartments by LP1/2 or muscle and further separated the layers by section: stricture, inflamed noninflamed and normal. Then Cellchat, an algorithm for analyzing cell-cell communication at the single-cell level[3]was used for inferring and quantifying the cell-cell communication within different sections. Kegg pathway enrichment analysis was performed using fgsea, fast gene set enrichment analysis: minSize = 10, maxSize = 500, nperm=100000. Genesets c2.cp.kegg.v2023.1.Hs.symbols.gmt was used in the analysis and these gene sets were obtained from MSigDB. Pathways that were significant (FDR<0.05) in one cell type or more are shown.

*Cyclic immunofluorescence staining and analysis*

Cyclic multiplex immunofluorescence (cycIF) was performed as described in Molina et al. using the antibodies described in Supplemental Table 2[4]. Briefly, formalin-fixed, paraffin-embedded (FFPE) sections of human full thickness resection specimens were deparaffinized, rehydrated and subjected to heat-induced epitope retrieval. After blocking with normal donkey serum slides were stained with primary antibodies overnight. Unbound antibodies were removed by washing and slides were stained with fluorescently conjugated secondary antibodies made in donkeys (Supplemental Table 2). Slides were again washed, DNA stained with DAPI (Invitrogen) and mounted in ProLong Diamond Antifade Mountant with DAPI (Invitrogen). Stained slides were imaged using an Axio Scan (Zeiss). Following each round of imaging, coverslips were removed and antibodies stripped. Acquired images were first subjected to shading correction, as well as stitching using the Zeiss ZEN blue DESK (version 3.0) image processing software. The alignment across subsequent rounds was performed on their respective nuclear (DAPI) channel via FIJI and applying the transformation to the remaining channels[5, 6]. The multi-channel image pyramid was generated using Bioformats2Raw and Raw2OMETiff programs (both Glencoe Software, Seattle, WA) and finally loaded into Visiopharm (Visiopharm A/S, Denmark) for analyses. A representative 2 mm x 2 mm ROI was centered on the tissue, where segmentation and heatmap APPs were applied for individual markers. All APPs were run with the same parameters across the experimental groups within the same tissue type to ensure data integrity. Numerical values were extracted and processed using Microsoft Excel and R and converted to a FlowJo-compatible format (FCS) for further analyses.

*Quantitative reverse transcriptase polymerase chain reaction*

Gene expression levels of human *CHD11* and mouse *TNF* and *IL6* were determined via quantitative RT-PCR. Total RNA was isolated as described previously (RNAEasy Miniprep kit, Qiagen),[7] and reverse transcription and quantitative PCR performed according to manufacture's instructions (Applied Biosystems, Foster City, CA). The products for all primer pairs were verified by sequencing and relative differences were calculated using the comparative threshold cycle method (ddCt) by normalizing to CT values of 18S or GAPDH (reference genes). Quantitative RT-PCR was performed on cDNA (synthesized with iScript cDNA Synthesis Kit, Biorad) with iQ Sybr Green Supermix (Biorad) and gene specific primers. GAPDH or 18S were used as reference genes and the Pfaffl method was used to calculate fold changes in treated versus untreated samples.[7] The primer pairs used for gene expression analysis are summarized in Supplemental Table 3.

*Immunostaining and quantification*

For CDH11 staining optimal cutting temperature (OCT) embedded tissues were used. Fresh frozen tissues were sectioned, and immunohistochemistry staining was performed manually. The slides were blocked with 3% horse serum (Media preparation core, Cleveland Clinic, Cleveland, OH). As primary antibody, human CDH11 (clone: 667039, MAB17901, 1:500 dilution, R&D Systems, Minneapolis, MN) was added to each section on the slides, and incubated for 1 hour at room temperature. CDH11 was visualized with ImmPRESS HRP anti-mouse IgG polymer reagent kit (MP-7402, Vector Laboratories Inc, Newark, CA) in conjunction with the ImmPACT DAB detection kit (SK-4105, Vector Laboratories, Inc. Newark, CA). The slides were counterstained with Hematoxylin QS (H3404, Vector Laboratories Inc, Newark, CA).

For mouse fibronectin 1 (FN1) and collagen I (ColI) staining, briefly, formalin fixed paraffin embedded (FFPE) slides were deparaffinized using ClearRite, Flex 100, Flex 95 and water and then incubated with antigen retrieval buffer (DAKO, pH=6.0) at 95°C for 30 min. Slides were placed at room temperature for 20 minutes, washed twice with PBS, dried and outlined with Pappen around samples. Slides were blocked with 5% fetal bovine serum (FBS, Life Technologies Corporation, Grand Island, NY) for 30 minutes and then stained with primary antibodies including anti-FN1(Cat# ab268020) and anti-Col1 (Cat# ab34710, both Abcam, Waltham, MA, diluted 1:100) in 5% FBS. One section was used in parallel as a secondary control. Slides were left in the dark for overnight at 4°C. Next day slides were washed twice with PBS and incubated with secondary antibody (Goat anti-rabbit 488 or anti-rabbit 594, 1:500 in 5% FBS on all sections) for an hour. After washing three times with PBS, samples were incubated with 0.1% Sudan Black B in 70% ethanol for 30 minutes at room temperature and then were covered with vectashield mounting media with DAPI.

All slides stained with CDH11, FN1 and Col1 were scanned and acquired using Aperio Image Scope software (Leica Biosystems, IL). CDH11, FN1 and Col1 stained images were quantified using ImageJ software (Bethesda, MD). Starting with a non-diseased control slide, two different areas were marked by an investigator blinded to the disease diagnosis in submucosal and muscle layers and threshold was set. All other diseased slides were quantified using the same set threshold and integrated density was measured using ImageJ and graphs plotted using GraphPad Prism software (version 9.3.1, Boston MA).

*Immunofluorescence labeling and confocal microscopy*

Immunofluorescence labeling was performed as previously described.[2] Briefly, fresh snap frozen OCT-embedded intestinal sections of histologically non-IBD controls and UC, CD intestinal tissue were stored at -80°C prior to use. Frozen human and mouse tissue sections were cut at 8 μm thickness and mounted to glass slides, thawed at room temperature for 1 hour, and fixed in ice cold methanol for 10 minutes. Slides were then washed 3 times with PBS and blocked with 5% FBS for 1 hour to eliminate non-specific binding. Co-staining of CDH11 was performed by incubation with primary antibodies to CDH11 (R&D, Minneapolis, MN), with below antibodies respectively: α-smooth muscle at 1:200 dilution, vimentin at 1:1000 dilution, E-cadherin at 1:100 dilution, desmin at 1:200 dilution, CD31 at 1:20 dilution, CD45 at 1:400 dilution, CD68 at 1:200 dilution at 4°C overnight. Slides were rinsed three times with PBS and incubated with AlexaFluor 488 or 594 conjugated donkey-anti rabbit and donkey-anti mouse secondary antibodies (1:500) for 1 hour at 37ºC. For nuclear counterstaining Vectashield mounting medium with DAPI (Vector Laboratories) was used. Immunolabeled tissue sections were examined using Leica confocal microscope (Leica TCS-SP8-AOBS inverted confocal microscope, Leica Microsystems, GmbH, Wetzlar, Germany) and upright fluorescent microscope (Leica DM6B upright microscope with Leica DMC4500, Leica Microsystems, GmbH, Wetzlar, Germany) LAS X software in the Lerner Research Institute Microscopy and Tissue Imaging Core, Image analysis was performed using ImagePro and Image J software (NIH, Bethesda, MD, USA).

*RNA extraction from FFPE tissue*

RNA was isolated from 2 x 20 μm FFPE tissue sections using the AllPrep DNA/RNA FFPE kit (Qiagen, Germantown MD). Manufacturer's instructions were followed for RNA extraction with the exception that tissue curls were prepared with 1280 μL deparaffinization solution (Qiagen) and RNA was digested using 300 μL Buffer PKD + 20 μL Proteinase K for 25 mins at 56C. RNA was eluted in 15 μL of RNAase-free water. RNA concentration was determined using Tecan-1 Spark™10M plate reader NanoQuant plate (Tecan, Baldwin Park, CA).

*Gene Expression Analysis using NanoString nCounter platform*

Gene expression was analyzed using nCounter PlexSet Reagents, nCounter MAX/FLEX System and nSolver Analysis Software 4.0 (NanoString Technologies, Seattle, WA). Briefly, 150ng of RNA from each sample was hybridized with a pair of target specific oligonucleotide probes for each targeted gene, using a predesigned nCounter PlexSet-24 Reagent Pack. Samples were incubated at 67 °C for 16 hours, multiplexed, and loaded using the nCounter Prep Station and scanned on nCounter Digital Analyzer. Following quality control checks, mRNA counts for test genes were normalized in two steps: raw NanoString counts were first background adjusted with a Truncated Poisson correction using internal negative controls followed by technical normalization using internal positive controls. Data were then corrected for input amount variation through a Sigmoid shrunken slope normalization step using the mean expression of housekeeping genes (*HPRT1*, *GUSB*, *RPLP0*). A transcript was designated as below detection level if the raw count was below the average of the 8 internal negative control raw counts plus 2 standard deviations.

*Isolation of primary human intestinal cells*

For validation experiments, we grouped CDi and CDni together, termed CD non stricturing or CDns. This was done as the CDni portions of the resection are very small in size and not amendable to cell isolation. Surgical resection specimens were used for isolation of primary human intestinal cells, including human intestinal myofibroblasts (HIMF), human intestinal muscularis propria muscle cells (HIMC), human intestinal microvascular endothelial cells (HIMEC), lamina propria mononuclear cells (LPMC) and epithelial cells as previously described.[1, 2] HIMF were obtained as explants of surgically resected intestinal mucosa, grown to subconfluence in Dulbecco's minimal essential medium supplemented with 10% fetal bovine serum (FBS) and antibiotics, and established as long-term cultures fed twice weekly and subcultured at confluence.[1, 2] Primary HIMC were obtained from surgically resected intestinal specimen and isolated based on a previously described method used for esophageal muscularis propria muscle cells.[1, 8] Human intestinal epithelial cells were obtained upon dissection of mucosal strips, and removal of epithelial cells through ethylenediaminetetraacetic acid (EDTA) solution. Epithelial cells were collected by centrifuging the EDTA solution at 1500 rpm, and then lysed for further immunoblotting.[9] Isolation of HIMEC was performed as previously reported.[7, 10] This consisted of enzymatic digestion of intestinal mucosal strips followed by cell sorting of CD31-positive using a BD FACS Aria machine (BD, San Jose, CA). LPMC were isolated from CD and non-IBD control tissues as previously described.[7, 11]

*Cell cultures*

HIMF monolayers were cultured in 6-well plates (Corning, Corning, NY) at 150,000 cells/well for immunoblotting analysis. In all experiments HIMF were serum-deprived 24h prior to harvest, and all experiments were performed under serum-free conditions.

*Immunoblotting*

Protein extraction was performed using a RIPA lysis buffer containing 50 mM TRIS pH 7.5, 150 mM NaCl, 1% Triton X-100, 0.1% SDS, 1% Na-deoxycholate and 1% protease and phosphatase inhibitor cocktail (Sigma) as previously described.[1, 2] The concentration of proteins in each lysate was measured using the Bio-Rad protein assay (Bio-Rad Laboratories, Hercules, CA) according to manufacturer's recommendations. Immunoblotting was performed as previously described.[2] Equivalent amounts of proteins (10 µg) were separated using SDS-PAGE on a 10% Tris-glycine gel and transferred to a PVDF membrane (Millipore, Billerica, MA). Nonspecific binding was blocked by incubation with 5% milk or 5% BSA in 0.1% Tween 20/Tris-buffered saline (Fisher Scientific) for 30 min., followed by overnight incubation at 4°C with the primary antibody(s). The following antibodies were used: Fibronectin at 1: 2000 (Cat# 610077, BD, San Jose, CA); CDH11 at 1:1000 (Cat# 32-1700, Thermo Scientific, Lafayette, CO) and GAPDH at 1:2000 (Cat# 2275-pc-100, Trevigen, Gaithersburg, MD). Membranes were washed 6 times with 0.1% Tween 20/Tris-buffered saline, incubated with the appropriate horseradish peroxidase-conjugated secondary antibody (Sigma-Aldrich, Inc., St Louis, MO), washed again, and incubated with the chemiluminescent substrate (Super Signal; Pierce, Rockford, IL) for 5 minutes, after which they were exposed to film (Kodak).

*RNA interference*

HIMF were transfected with CDH11-specific small interfering RNA (siRNA) and their respective scrambled siRNA (Horizon Discovery, Cambridge, United Kingdom) using Lipofectamine 2000 (Thermo Scientific, Lafayette, CO). The siRNAs are pooled nucleotides with different sequences for each target in the same solution. After optimization of the experimental conditions, HIMF (1.5 x $10^5$) were plated in 6-well plates and allowed to attach overnight. Cell culture medium was removed and replaced with opti-MEM as well as lipofectamine and siRNA mixture. After transfection for 6 hours, the media was replaced with regular HIMF culture media and the cells were harvested at 24 hours, 48 hours and 72 hours post-transfection.

*Proteomics analysis of conditioned medium from human intestinal myofibroblasts*

HIMF were exposed to CHD11 activator (CDH11Fc) or PBS (control) for 48 h (nine replicates each), and conditioned media collected and analyzed by proteomics as described previously.[12] Briefly, samples were filtered with aa Amicon Ultra 3K molecular weight 0.5 mL centrifugal filter (Millipore # UFC500396) and then dried down in Speed-Vac before being reconstituted in 25 µL of 6 M Urea/Tris buffer. Samples were reduced with DTT, alkylated with iodoacetamide, and precipitated with cold acetone overnight at -20°C. Proteins were then digested in solution with trypsin (Promega # V5111) in 50 mM TEAB overnight at 37°C. A second aliquot of trypsin was added and digestion continued for 4 hours. Digestion was terminated by adding trifluoroacetic acid to a final concentration of 0.5-1%, and samples were desalted using PepClean C-18 spin columns (Thermo Scientific #89870). After clean-up, samples were reconstituted in 1% formic acid to a final volume of ~30 µL, for LC-MS analysis. The LC-MS system was a ThermoScientific Exploris 480 mass spectrometer equipped with a Vanquish Neo uHPLC system. The HPLC column was a 50 cm x 75 µm id Easy Spray Pepmap NEO C18, 2µm, 100 Å reversed- phase capillary chromatography column. Sample extracts (5 µL each) were injected and the peptides eluted from the column by an acetonitrile/0.1% formic acid gradient at a flow rate of 0.3 µL/min. The digest was analyzed using the data dependent multitask capability of the instrument acquiring full scan mass spectra to determine peptide molecular weights and product ion spectra to determine amino acid sequence in successive instrument scans. Exploris data were analyzed by using all CID spectra collected in the experiment to search the human SwissProt database using the program Proteome Discoverer 2.4 (protein FDR rate was set to 1%), and obtain the label free quantitation (LFQ) intensities. The LFQ intensity values were normalized and used for downstream analysis (differential abundance, volcano plot, pathway analyses) using the *R* package *DEP*.[13]

*Next generation RNA sequencing*

Total RNA was extracted from $5x10^5$ non-IBD HIMF treated each either with scrambled siRNA only or *CDH11* siRNA in 4 separate experiments using the RNeasy Mini Kit (Qiagen, Germany). RNA quality was assessed using the Agilent bioanalyzer and samples with RNA quality (RIN) score of >9.0 were used in RNA-sequencing. RNA concentrations were determined using Qubit® 3.0 Fluorometer (Invitrogen, Life Technologies). RNA-sequencing libraries were generated using Illumina TruSEQ kits following the manufacturer's protocol and libraries were sequenced using an Illumina NovaSeq 6000 following Illumina reagents and protocols. Paired-ended 100 base pair reads were trimmed and checked for quality with FastQC,[14] and aligned with human genome (GRCh38.p13) using *Subread* package.[15] Overall, an average of 94% of reads aligned uniquely. Gene counts were determined by the number of uniquely aligned, unambiguous reads (Subread: featureCounts, v1.5.2) and annotated (GRCH38, Ensembl 99 release). On average, 70% of reads in each sample were successfully assigned. Raw counts were loaded into R (ver. 4.2.1)[16] and filtered to exclude transcripts that were lowly expressed (counts per million reads mapped [CPM] < 1) before performing differential expression (DE) analysis with *limma*[17] and *edgerR*.[18] The resulting count values were used for library size normalization, log2 transformation and downstream analyses. Heatmaps and volcano plots were generated using *pheatmap*[19] and *EnhancedVolcano* packages[20], respectively. Differentially expressed genes (DEGs) with Benjamini-Hochberg adjusted p-value < 0.05 and Log2FoldChange >1 or <-1 were considered statistically significant. Functional analysis of significant DEGs including KEGG enrichment were performed using R packages *clusterProfiler*[21, 22] and *gprofiler*.[23] Additional graphs were generated using the *tidyverse* set of packages.[24] The *R* package *grateful* was used to generate a summary list of all packages used in the study.[25] RNA sequencing data have been deposited in Gene Expression Omnibus (GEO).

*Antibodies*

Antibodies were purchased from the following companies: α-smooth muscle actin (α-SMA) (catalog ab5694), vimentin (catalog ab45939); E-cadherin (catalog ab76055), desmin (catalog ab15200), CD31(catalog ab28364), CD45 (catalog ab10558), CD68 (catalog ab125212), fibronectin for immunolabeling (catalog ab32419) (all from Abcam, Cambridge, MA). CDH11 for immunolabeling (catalog MAB1790, R&D, Minneapolis, MN), CDH11 for immunoblotting (catalog 32-1700, Thermo Scientific, Lafayette, CO), fibronectin for immunoblotting (catalog 610077, BD, San Jose, CA), GAPDH (catalog 2275-pc-100, Trevigen, Gaithersburg, MD).

*Animals*

CDH11 KO mice with CD1/B6/129 background and for the CDH11 blocking experiments wild type Balb/c mice were purchased from Jackson Laboratory (Bar Harbor, ME). All mouse lines were bred, genotyped and maintained in the Biomedical Research Unit at the Cleveland Clinic using standard protocols. Genotyping was performed by toe clipping or ear notching. Littermate controls were used for experiments. All studies were approved by the Cleveland Clinic Institutional Animal Care and Use Committee (IACUC protocols 2018-2023 and 2019-2137).

*Dextran sodium-sulfate induced colitis*

All animal studies were conducted following protocol approved by the Cleveland Clinic

Institutional Animal Care and Use Committee (IACUC protocols 2018-2023 and 2019-2137), and followed ARRIVE guidelines.

**ARRIVE ESSENTIAL 10 CHECKLIST**

**Study 1. Effect of CDH11 blockade on experimental colitis**

| | |
|---|---|
| Study design | Male BALB/C mice (6-8 weeks old) with or without 3% DSS, treated with blocking monoclonal antibody. Experimental groups received 0.5mg H1M1 through intraperitoneal injections every other day, while control group received mouse IgG2b isotype control. |
| Sample size | 10 mice per group |
| Inclusion and Exclusion Criteria | Male mice were included in the study |
| Randomization | Mice were randomly divided into two groups: isotype control or H1M1 group |
| Blinding | An IBD pathologist scored the tissues in a blinded fashion for inflammation and fibrosis using H&E sections. Fibrosis was also assessed by Sirius red staining, Col1 or FN immunolabeling and quantified using integrated density measurements for the submucosa and the muscularis propria separately |
| Outcome measures | Improvement in fibrosis, defined as reduction in fibrosis area in the submucosa of the H1M1 group. |
| Statistical methods | We evaluated intestines of H1M1-treated versus isotype- |

| | |
|---|---|
| | treated mice to see if a difference is observed when fibrosis initiates. |
| Experimental animals | Wild type Balb/c mice were purchased from Jackson Laboratory (Bar Harbor, ME). All mouse lines were maintained in the Biomedical Research Unit at the Cleveland Clinic using standard protocols. |
| Experimental procedures | Wildtype (WT) mice were subjected to two cycles of dextran-sodium sulfate (DSS) administration (3%) followed by post-DSS recovery. Cdh11 blocking antibody H1M1 or isotype antibody were given daily starting day 1 of the experiment. Endpoint was day 44. Animals were euthanized by $CO_2$ asphyxiation followed by cervical dislocation. Clinical disease activity was determined every other day by measuring body weight, stool consistency and presence of occult or overt blood in stool. Severity of DSS induced colitis was evaluated by measuring the body weight loss and calculating the clinical score. Inflammation score was determined using H&E sections by an IBD pathologist in a blinded fashion. Severity of fibrosis was evaluated through picrosirius red staining, Col1 or FN immunolabeling and quantified using integrated density measurements for the submucosa and the muscularis propria separately. Thickness of the intestinal layers was measured for mucosa, submucosa |

| | |
|---|---|
| | and muscularis propria. Gene expression of TNF and IL6 in intestinal tissues was determined using q-RTPCR. |
| Results | CDH11 blockade reduced fibrosis and inflammation in mice. |

**Study 2. Effect of CDH11 knockout on experimental colitis**

| | |
|---|---|
| Study design | Effect of CDH11 knockout on experimental colitis |
| Sample size | 12 mice per group |
| Inclusion and Exclusion Criteria | Male mice were included in the study |
| Randomization | Knockout and wild type mice were used. |
| Blinding | An IBD pathologist scored the tissues in a blinded fashion for inflammation and fibrosis using H&E sections. Fibrosis was also assessed by Sirius red staining, Col1 or FN immunolabeling and quantified using integrated density measurements for the submucosa and the muscularis propria separately |
| Outcome measures | Improvement in fibrosis (defined as reduction of the fibrosis area in the submucosa of the CDH11 KO DSS group and no fibrosis in the non−DSS treatment). |
| Statistical methods | We evaluated intestines of CDH11 KO versus wildtype mice to see if a difference is observed when fibrosis initiates. |
| Experimental animals | CDH11 knockout (KO) mice with B6/129 background were kindly provided by Jackson (Bar Harbon, USA).<br>All mouse lines were bred, genotyped and maintained in the |

| | |
|---|---|
| | Biomedical Research Unit at the Cleveland Clinic using standard protocols. Genotyping was performed by toe clipping or ear notching. Littermate controls were used for experiments. |
| Experimental procedures | CDH11 null (KO) mice and wildtype (WT) littermates were subjected to two cycles of dextran-sodium sulfate (DSS) administration (2%) followed by post-DSS recovery. Endpoint was day 44. Animals were euthanized by $CO_2$ asphyxiation followed by cervical dislocation. Clinical disease activity was determined every other day by measuring body weight, stool consistency and presence of occult or overt blood in stool. Severity of DSS-induced colitis was evaluated by measuring the body weight loss and calculating the clinical score. Colonic sections were fixed and stained with hematoxylin and eosin (H&E) for histological examination. Thickness of the intestinal layers was measured for mucosa, submucosa and muscularis propria. Gene expression of TNF and IL6 in intestinal tissues was evaluated by q-RTPCR. |
| Results | CDH11 knockout mice exhibited reduced fibrosis. |

Chronic DSS colitis was induced as previously described.[2, 26, 27] After a preliminary dose-finding study, 2 % DSS (35–50 000 kDa; MP Biomedicals, Santa Ana, CA) in drinking water was chosen as optimal for the CD1/B6/129 CDH11 KO mouse strain and 3% for the Balb/c mouse strain. Chronic colitis was induced by DSS in drinking water for 8 days followed by a recovery period of 14 days with normal drinking water, and this was defined as one cycle of DSS. The DSS cycle was repeated twice. Control mice received normal drinking water throughout. Experimental colitis was performed in at least three independent experiments. All mice per group were age-matched and were co-housed as derived from the same litter and per genotype to minimize influence from differences in flora composition.[28] Clinical disease activity was determined every other day by measuring body weight, stool consistency and presence of occult or overt blood in the stools as previously described. Endpoint was day 44. Animals were euthanized by $CO_2$ asphyxiation followed by cervical dislocation.

*Chronic DSS model with CDH11 blockade in BALB/c WT mice*

6-8-week old male BALB/c mice (Jackson laboratories, Bar Harbor, ME) were used for CDH11 blockade experiment. Chronic colitis was induced by DSS for 8 days followed by a recovery period of 14 days with normal drinking water, and this was defined as one cycle of DSS. The DSS cycle was repeated twice. Control mice received normal drinking water throughout. Experimental groups received 0.5mg H1M1 through intraperitoneal injections every other day, while control group received mouse IgG2b isotype control (Bio X Cell, NH, USA). The optimal dose for the antibody was determined as described in *Results*. The starting point of antibody injections was on Day 0. Clinical disease activity was determined every other day as previously described.[2, 29, 30] Body weights, stool consistency and occult blood or the presence of gross blood per rectum were recorded every other day. Two investigators blinded to the protocol independently assessed the clinical score as previously described.[30] Briefly, weight loss of 1–5%, 5–10%, 10–20%, and >20%

was scored as 1, 2, 3, and 4, respectively. For stool consistency, 0 was scored for well-formed pellets, 2 for pasty and semiformed stools, which did not stick to the anus, and 4 for liquid stools that remained adhesive to the anus. Bleeding was scored 0 for no blood in hemoccult, 2 for positive hemoccult, and 4 for gross bleeding from the rectum. Weight, stool consistency, and bleeding sub-scores were added and divided by 3, resulting in a total clinical score ranging from 0 (healthy) to 4 (maximal activity of colitis). Animals were euthanized by $CO_2$ asphyxiation followed by cervical dislocation.

*Experimental fibrosis endpoints*

At the end of the experiment the entire colon was removed, cleaned and measured from the ileocecal junction to the anus. Tissue was procured from the descending colon. Histology was performed on paraffin embedded, 3 μm-thick transverse sections stained with hematoxylin/eosin or picrosirius red. Slides were scored by an experienced pathologist (I.O.G.) blinded to the experimental groups using an inflammation score as previously described.[2, 31] Briefly, inflammation scoring was performed using hematoxylin & eosin (H&E) slides based on inflammation infiltration (0-3), extent of inflammation (0-3), crypt damage (0-4) and percentage of involved area (0-4).

For quantitation of picrosirius red, Col1 and FN1 stained slides (details of stains and staining procedure described above), images were scanned and acquired using Aperio ImageScope software (Lieca Biosystems, IL). Acquired images were quantified using ImageJ software (Bethesda, MD). Starting with a non-diseased control slide, threshold was set using submucosal and muscle layers of a no DSS wildtype control animal. The set threshold was used to quantify diseased animal slides and integrated density was measured using ImageJ and graphs plotted using

GraphPad Prism software (version 9.3.1, Boston, MA). The thickness of the intestinal wall layers including muscularis mucosa, submucosa and muscularis propria were calculated as the mean value of different areas marked on each layer per mouse on well oriented cross sections using Aperio ImageScope software (Leica Biosystems, IL) and graphs were plotted using GraphPad Prism (version 9.3.1, Boston, MA).

*Statistical analysis*

Data were analyzed using analysis of variance (ANOVA) for independent groups. Repeated measures for the same experiment were analyzed by using Student's paired t test. Values were expressed as mean ± SEM, and statistical significance was set at $p<0.05$. All analyses were performed using SAS (version 9.3; The SAS Institute Inc., Cary, NC) unless otherwise stated for the RNA sequencing results.

#### SUPPLEMENTAL RESULTS

*Major signaling pathways identified in the single cell RNA sequencing analysis across the entire dataset*

Major signaling pathways active in CD compared to NL using pseudobulk analysis were related to immune activation and host defense (Supplemental Figure 2A). Pathways modulated in CDi and CDs compared to CDni were an upregulation of toll like receptor signaling, JAK/STAT signaling, natural killer cells and cytokine/chemokine signaling. Of note an increase in ECM receptor interaction and focal adhesion was noted in most cell types. A dowregulation in CDi and

CDs compared to CDni was found in pathways linked with muscle contraction, fatty acid, glucose and amino acid metabolism and oxidative phosphorylation (Supplemental Figure 2A).

Overall and across all MP CD segments versus NL comparable pathways to the LP1/2 layers were found to be active, specifically pathways related to immune activation and host defense. As already noted in the pseudobulk analysis the changes between CD segments were small (Supplemental Figure 2B). Notable findings were an upregulation of cytokine/chemokine, PPAR and calcium signaling in CDs as well as a trend towards increased signaling of B cell receptors, FC receptor, T cell receptor signaling as well as JAK/STAT/WNT signaling and endocytosis (Supplemental Figure 2B).

*Major signaling pathways identified in the single cell RNA sequencing analysis in the stromal compartment*

On a per cell type pathway analysis the major stromal pathways being active in CD compared to NL were NOD like receptor signaling, chemokine and cytokine signaling, antigen processing and presentation and broad immune pathways. Stromal cells in CDi and CDs compared to normal and CDni upregulated ECM receptor interaction and muscle related pathways (Supplementary Figure 3A).

Comparable to the LP1/2 layer, on per cell type pathway analysis in the MP the pathways upregulated in CD compared to NL were broad immune activation, cytokine/chemokine and innate immune pathways. There was minimal difference between the CDni, CDi and CDs segments with the exception of an upregulation of focal adhesion and proliferation pathways (Supplementary Figure 3B).

*Stromal populations identified in previous publications and queried in our work*

In an effort to validate our findings, we examined how stromal populations identified in previously published works aligned with our data. A recent investigation by Kong et al.[32] identified two populations expressing HHIP/NPNT+ and GREM1/GREM2+, which we confirmed to be present in our data (HHIP/NPNT+ and DES+/ACTG2+, respectively). Interestingly we found that the HHIP/NPNT+ population is overrepresented in the LP1/2 layers while the DES/ACTG2+ population is overrepresented in the MP layer (Supplemental Figure 1B&C, Supplemental Figure 4A&B). We found a trend towards increased abundance of the DES/ACTG2 neighborhoods in CDs versus NL (LP1/2 layer), neighborhoods for neither population reach significance in our work. A different population previously implicated in disease has been inflammatory fibroblasts, first noted in single cell data by Smillie et al.[33]. Using the top markers from their work, we map the signature to our MMP/WNT5A+ fibroblast population which we note as specifically increased in abundance in CDs compared to NL, CDni, and CDi as well as in CDi versus NL (Supplemental Figure 4B). FAP has been implicated in multiple fibrotic diseases and may serve as a therapeutic target or biomarker[34-36]. We interrogated our dataset for FAP+ fibroblasts and found the highest expression to be present on our MMP/WNT5A+ and ECM$^{high}$ fibroblasts, both of which we see increased in abundance in CDs compared to CDi layer (Supplemental Figure 4A&B).

#### SUPPLEMENTAL FIGURES AND FIGURE LEGENDS

**Supplemental Figure 1. Quality metrics of the scRNAseq dataset and differences in cell abundance by tissue layer. A.** Violin plot of quality control metrics for each tissue sample of number of genes/cell (nFeature_RNA), unique UMIs/cell (nCount_RNA), and mitochondrial percentage per cell (percent.mt) colored by patient. **B.** Violin plot of mitochondrial percentage by cell type. **C&D.** Differential abundance beeswarm plots from LP1, LP2 and MP layers by cell type. Entire dataset in (**C**) and stromal compartment in (**D**). Each dot is a neighborhood of cells calculated using miloR. Neighborhoods that reach significance (spatial FDR < 0.1) are colored by log fold-change.

**Supplemental Figure 2. Altered transcriptome at the global level using pseudobulk analysis. A.** Heatmap of the fGSEA normalized enrichment scores (NES) in the LP1/2 layers of significantly enriched KEGG pathways in one or more cell types. An asterisk denotes statistical significance (FDR<0.05). The columns of the matrix represent cell types and the rows of the matrix indicate KEGG pathways. The column annotation indicates the contrast (stricture vs normal, inflamed vs normal and noninflamed vs normal). **B.** Heatmap of the fGSEA normalized enrichment score in the MP layer of significantly enriched KEGG pathways in one or more cell types. An asterisk denotes statistical significance (FDR<0.05). The columns of the matrix represent cell types and the rows of the matrix indicate KEGG pathways. The column annotation indicates the contrast (stricture vs normal, inflamed vs normal and noninflamed vs normal).

**Supplemental Figure 3. Altered transcriptome at the stromal level using pseudobulk analysis. A.** Heatmap of the fGSEA normalized enrichment score in the LP1/2 layer of significantly enriched KEGG pathways in one or more cell types. An asterisk denotes statistical significance (FDR<0.05). The columns of the matrix represent cell types and the rows of the matrix indicate KEGG pathways. The column annotation indicates the contrast (stricture vs normal, inflamed vs normal and noninflamed vs normal). **B.** Heatmap of the fGSEA normalized enrichment score in the MP layer of significantly enriched KEGG pathways in one or more cell types. An asterisk denotes statistical significance (FDR<0.05). The columns of the matrix represent cell types and the rows of the matrix indicate KEGG pathways. The column annotation indicates the contrast (stricture vs normal, inflamed vs normal and noninflamed vs normal).

Supplemental Figure 4. **Dataset annotation for previously described stromal cell populations associated with IBD. A.** Dot plot of canonical markers used to define previously described stromal cell types with relevance to IBD. Our own cell annotation is depicted on the left. Average gene expression is shown by color intensity and percent of cells expressing each gene is shown by size. **B.** Abundance of previously described cell populations by segment in our dataset. **C.** Dot plot of the major cell neighborhoods depicting *CDH11* expression. Average gene expression is shown by color intensity and percent of cells expressing each gene is shown by size.

Supplemental Figure 5. **Cell with cell interaction analysis.** Heatmaps of the incoming (**A & C**) and outgoing (**B &** D) interaction strength for each cell type by segment and separated into LP1/2 and MP layers. Interaction strength was calculated using cellChat[3].

Supplemental Figure 6. Full thickness sections of NL, normal; UC, ulcerative colitis; CDns, Crohn's disease, non-strictured; CDs, CD strictured patients were dual-immunolabeled for CDH11 (red) and either αSMA (**A**), vimentin (**B**), desmin (**C**), CD68 (**D**), E-cadherin (**E**), CD31 (**F**) or

CD45 (**G**) (green). Representative confocal images are shown. Arrows point to colocalization of CDH11 with markers. Stainings are representative for n=4 per group.

**Supplemental Figure 7. Matrisome proteins secreted by primary human intestinal myofibroblasts in response to cadherin-11 activation.** Each plot represents one core matrisome protein. The log2 fold change with 95% confidence interval is delineated (n=9 each group).

**Supplemental Figure 8. Global gene expression in human intestinal myofibroblasts in response to CDH11 knockdown. A.** Top 30 up (blue) and downregulated (red) genes upon *CDH11* knockdown. **A.** Heatmap correlating core matrisome genes with and without *CDH11* knockdown (KD). **B.** Significantly up (yellow) and downregulated (red) genes upon *CDH11* knockdown. **C.** GO category netplot analysis of clusters upon CDH11 knockdown (n=4 each group).

**Supplemental Figure 9**. H1M1 pharmacokinetics (PK) study in wild type BALB/c mice to establish the appropriate dose of antibody. H1M1 0.5 mg was administered intraperitoneally at different intervals and serum concentrations of the antibody measured at the delineated timepoints. A minimum plasma H1M1 level >200µg/ml is considered optimal for efficacy.[37, 38]

### Supplemental Figure 1

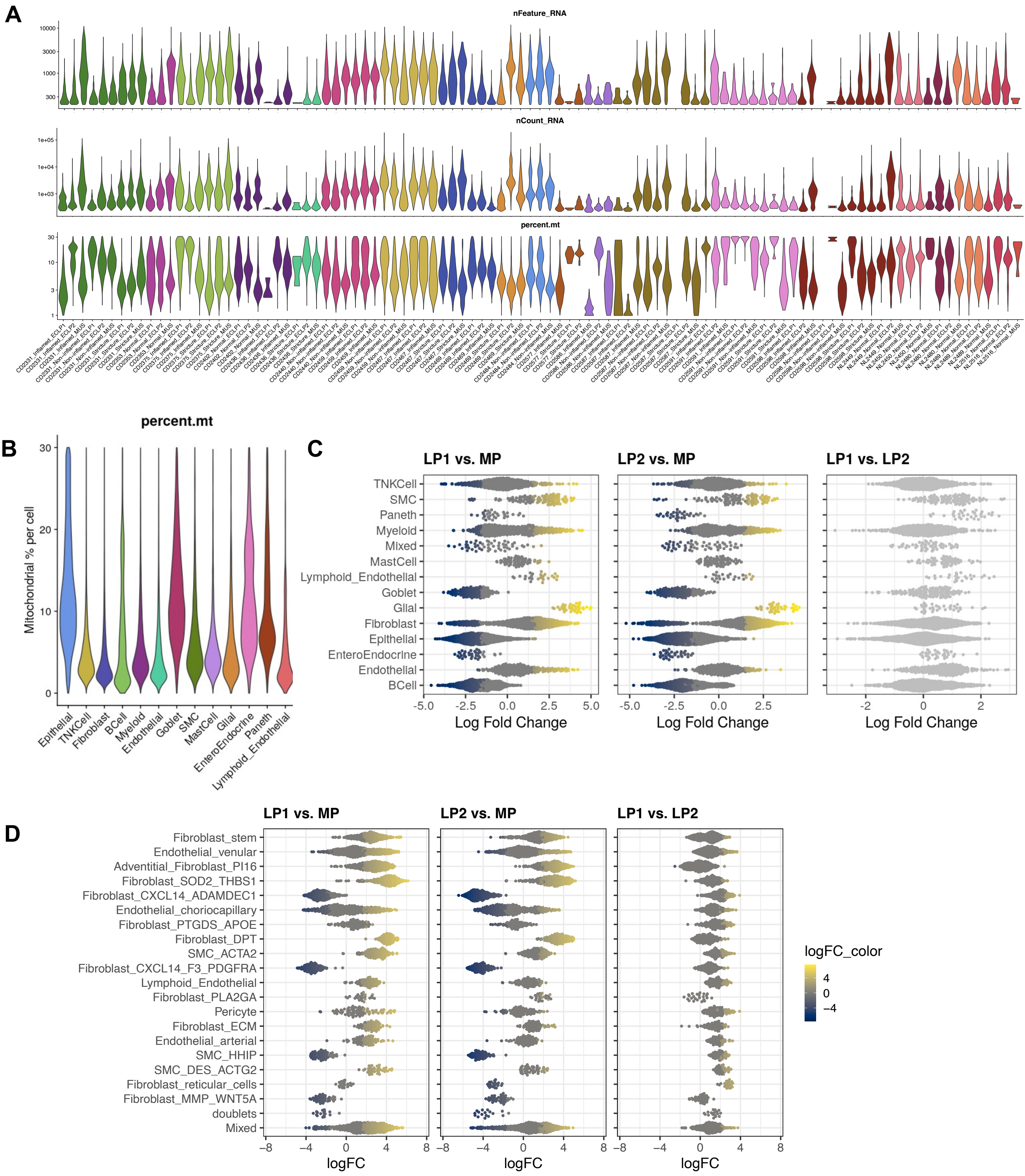

### Supplemental Figure 2

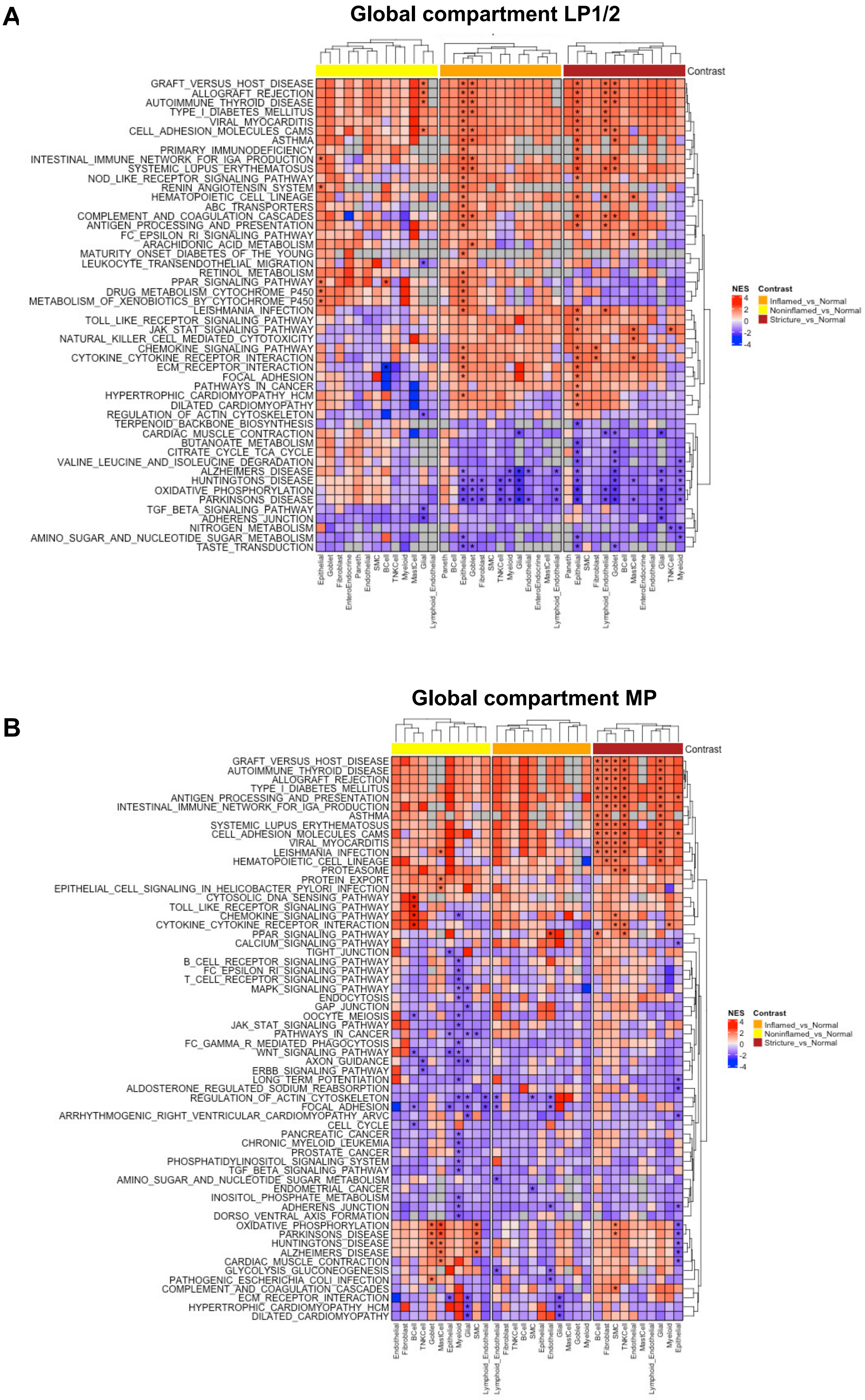

#### Supplemental Figure 3

A

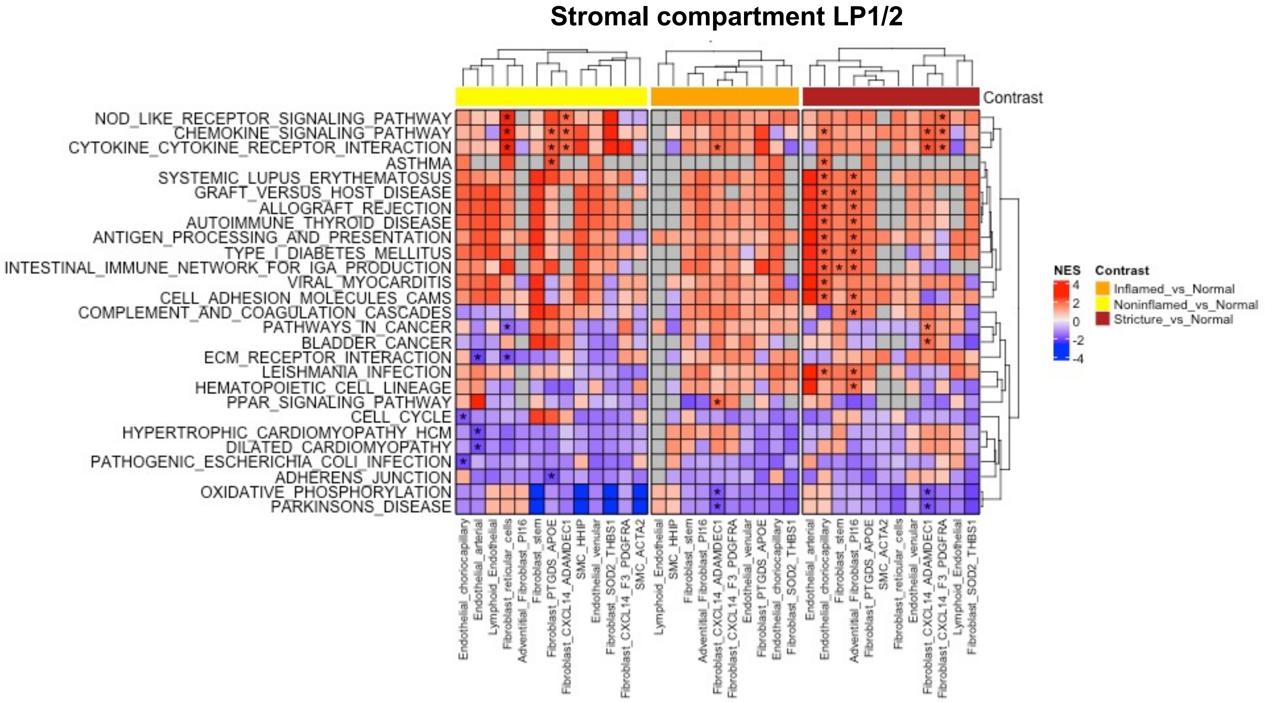


B

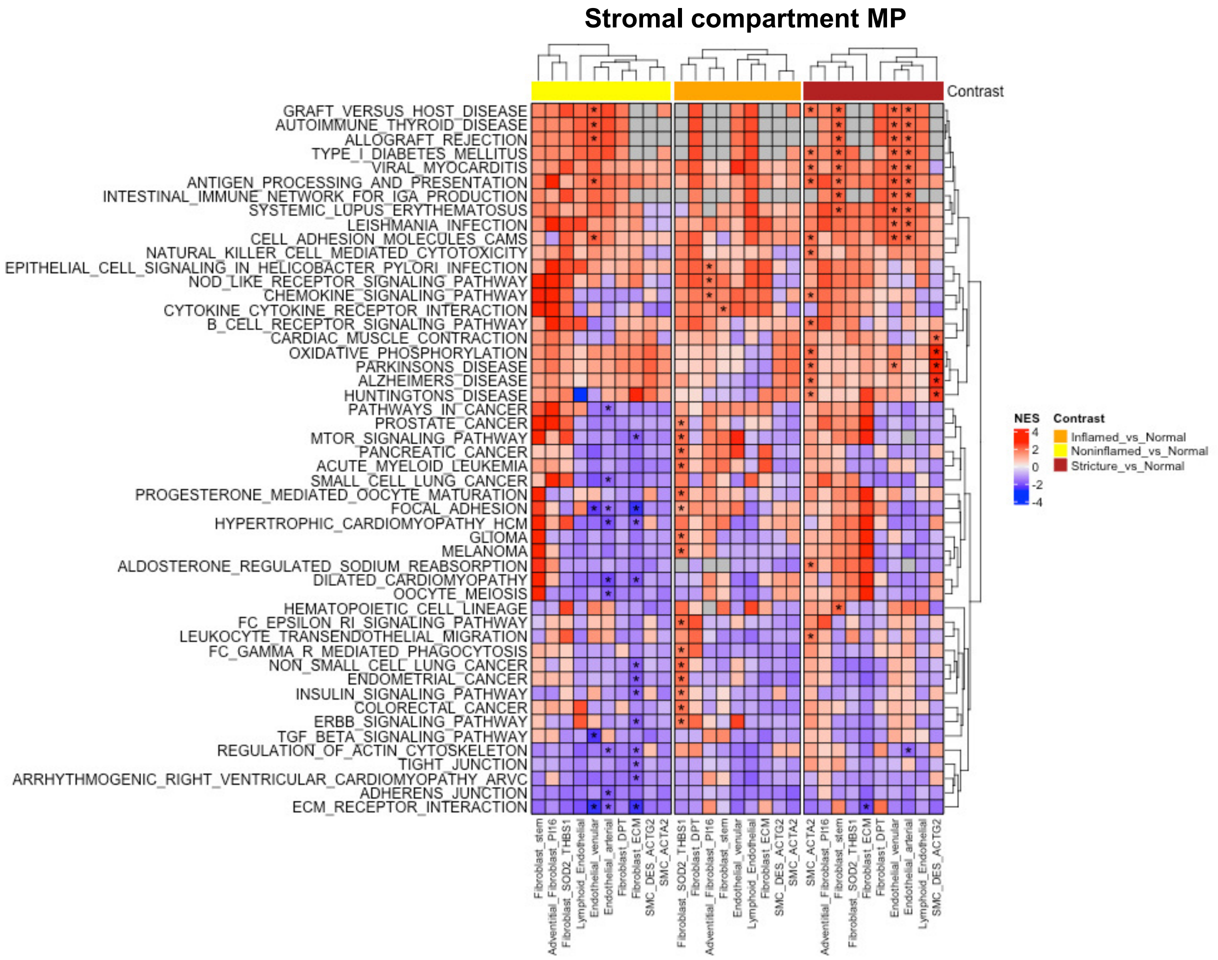

### Supplemental Figure 4

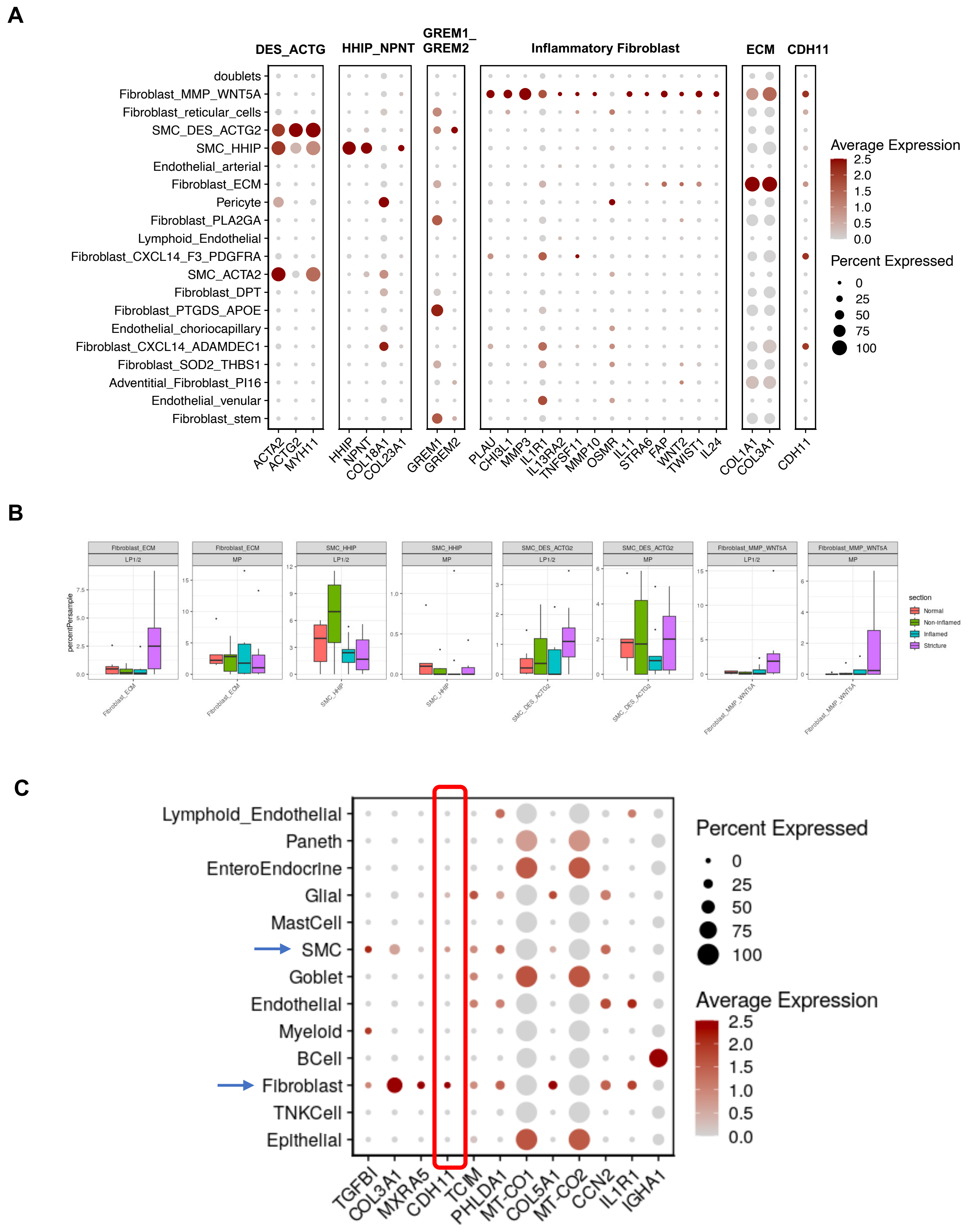

### Supplemental Figure 5

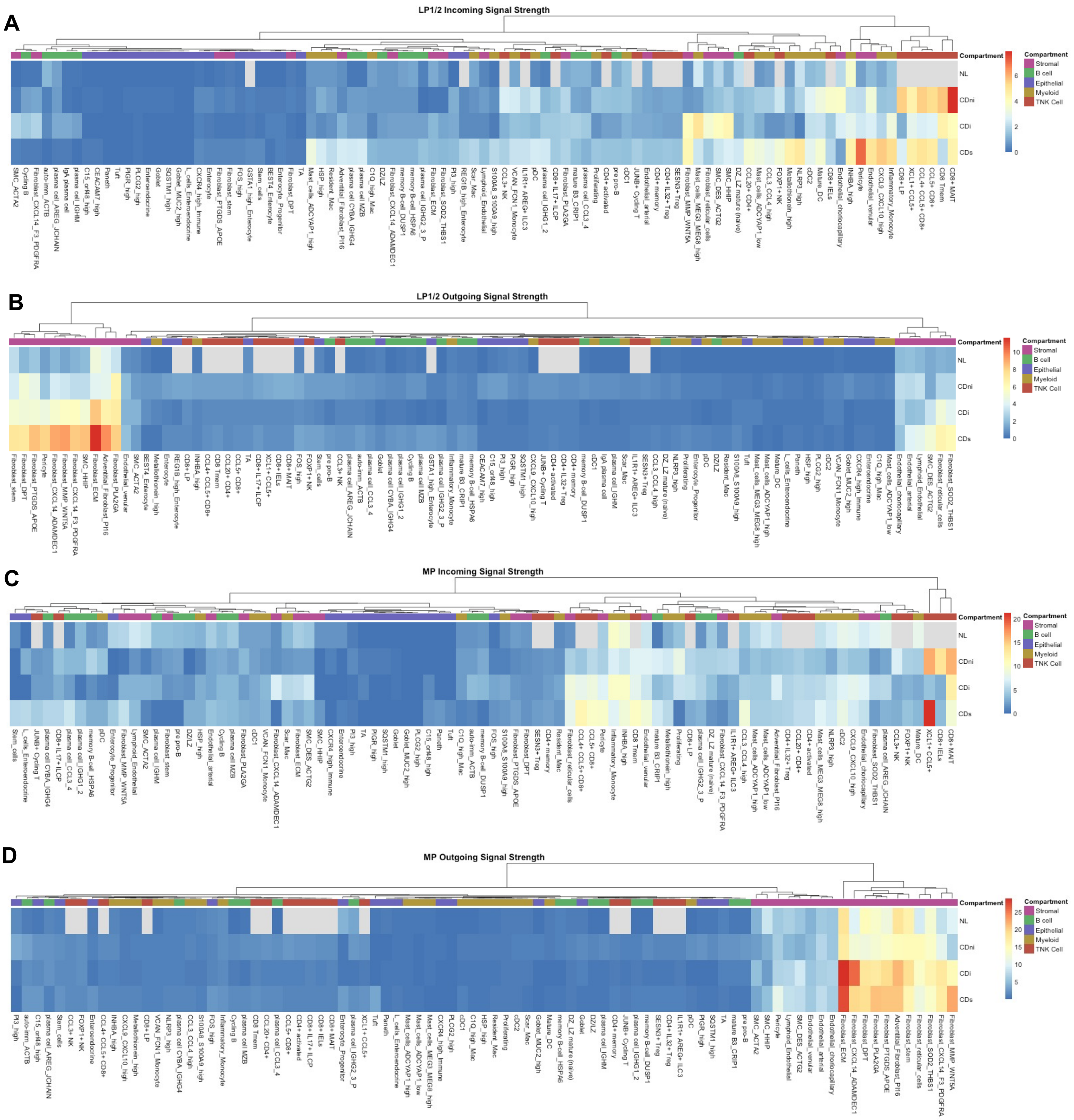

#### Supplemental Figure 6

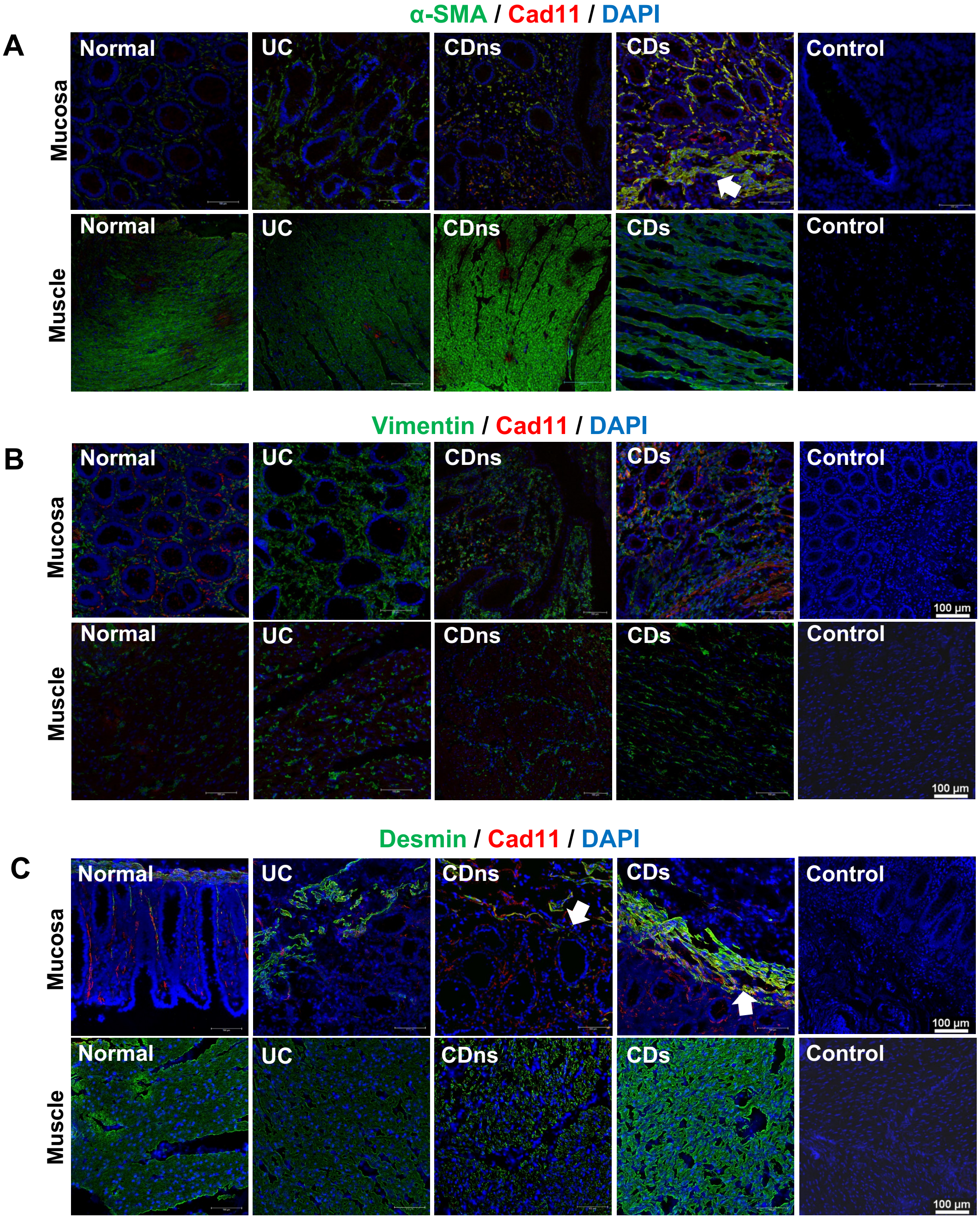

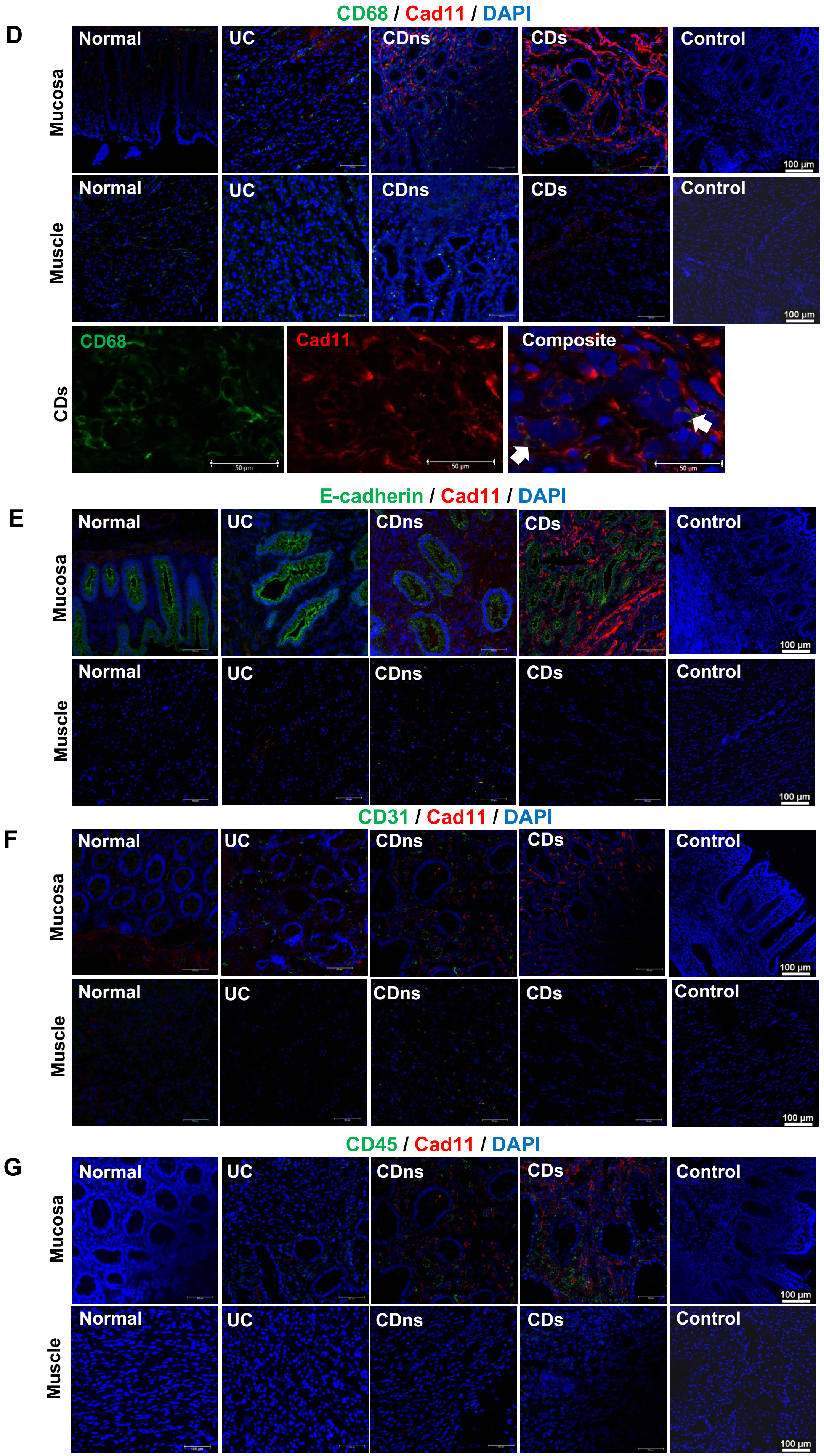


Supplemental Figure 6

**Supplemental Figure 7**

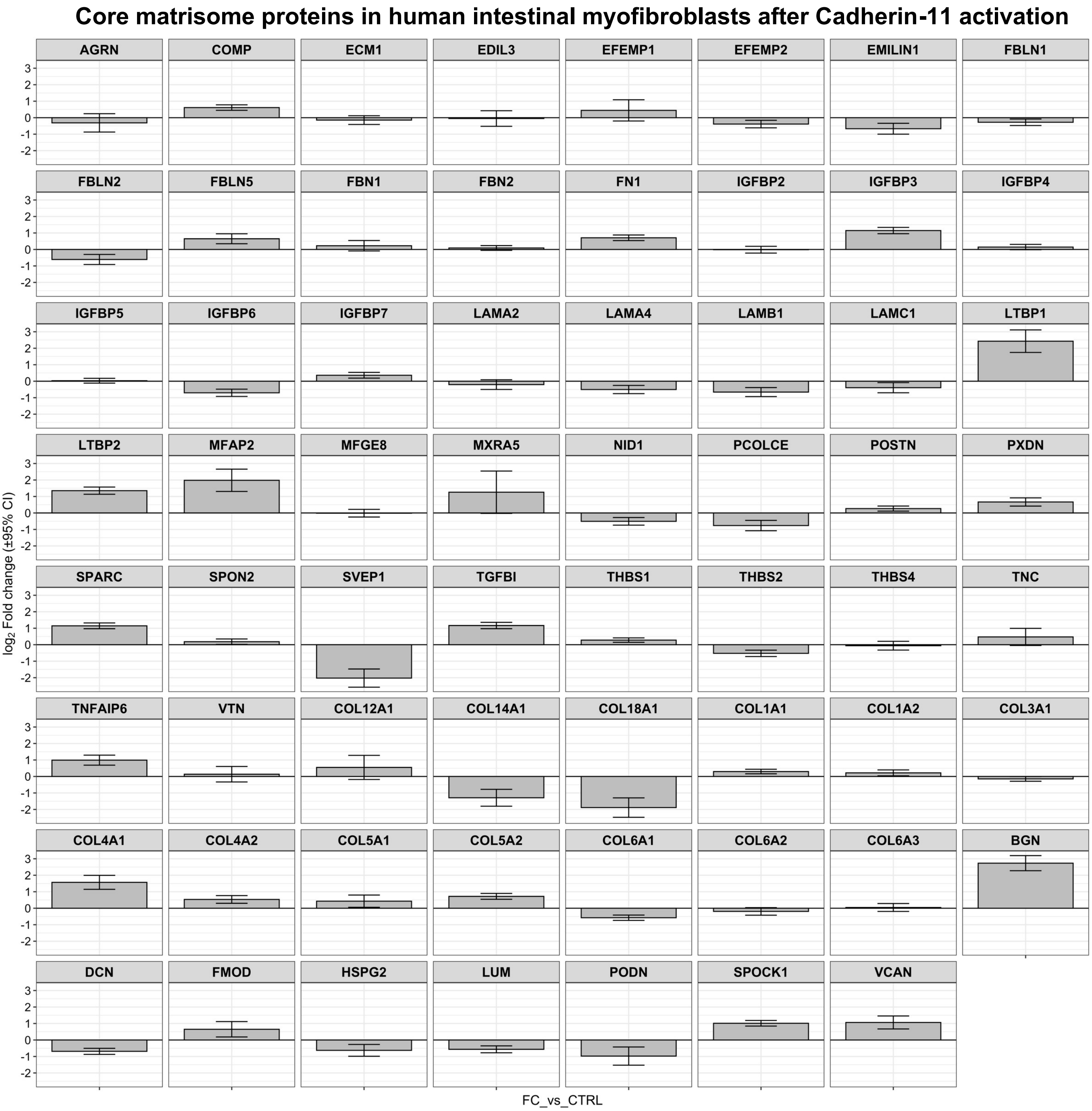

**Supplemental Figure 8**

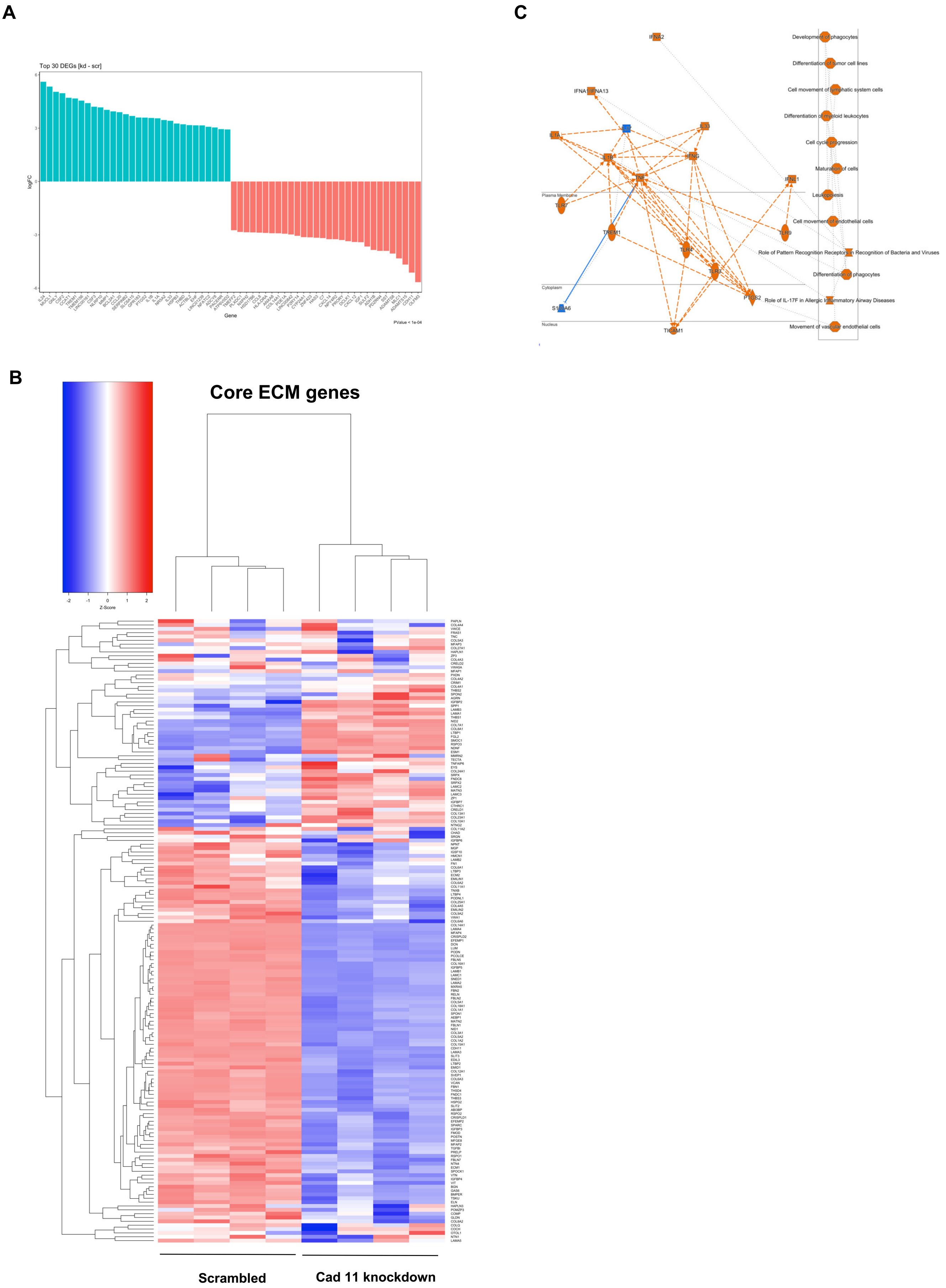

#### Supplemental Figure 9

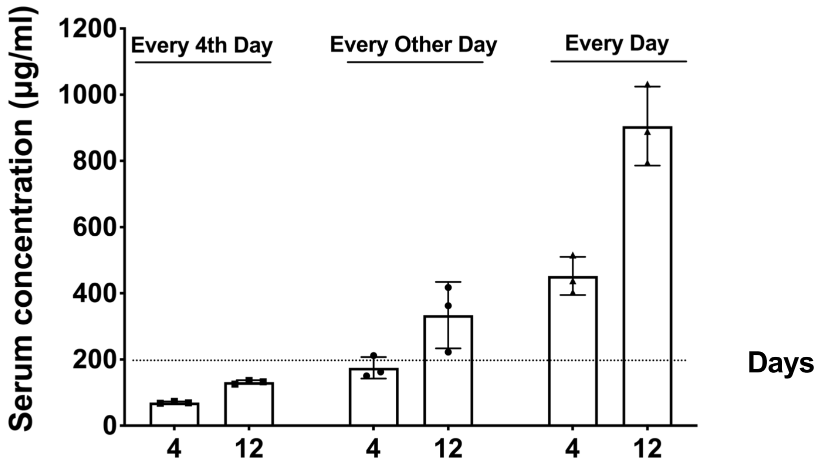
