## Supplemental Tables for "Stricturing Crohn’s disease single-cell RNA sequencing reveals fibroblast heterogeneity and intercellular interactions"

#### Supplemental Table 1. Chemistry type for each sample for 10x Chromium

Single cell suspensions were processed using either the 3’ v3.1 or 5’ v2 chemistries as described in *Methods*. Chemistry type for each sample is listed below and annotations for each sample correspond to the data uploaded into GEO.

| Sample ID | Diagnosis | Chemistry |
|---|---|---|
| CD2331 | CD | 10x 3’ v3.1 |
| CD2375 | CD | 10x 3’ v3.1 |
| CD2436 | CD | 10x 3’ v3.1 |
| CD2440 | CD | 10x 3’ v3.1 |
| CD2459 | CD | 10x 3’ v3.1 |
| CD2467 | CD | 10x 3’ v3.1 |
| CD2469 | CD | 10x 3’ v3.1 |
| CD2484 | CD | 10x 3’ v3.1 |
| CD2577 | CD | 10x 5’ v2 |
| CD2586 | CD | 10x 5’ v2 |
| CD2587 | CD | 10x 5’ v2 |
| CD2591 | CD | 10x 5’ v2 |
| CD2598 | CD | 10x 5’ v2 |
| NL2353 | NL | 10x 3’ v3.1 |
| NL2402 | NL | 10x 3’ v3.1 |
| NL2449 | NL | 10x 5’ v2 |
| NL2450 | NL | 10x 5’ v2 |
| NL2480 | NL | 10x 5’ v2 |
| NL2489 | NL | 10x 5’ v2 |
| NL2516 | NL | 10x 5’ v2 |

Abbreviations: CD, Crohn’s disease; NL, normal

**Supplemental Table 2. Antibodies used in cyclic immunofluorescence staining**

| Target protein | Product number | Company |
|---|---|---|
| Alpha SMA | Clone 1A4, A 2547, mouse monoclonal | Sigma |
| Desmin | PA5-16705, rabbit polyclonal | Thermo |
| Vimentin | Clone 28061, rat monoclonal | R&D |
| Cadherin 11 | Clone 667039, mouse monoclonal | R&D |

**Supplemental Table 3. Primers used for quantitative and qualitative polymerase chain reaction**

| Gene | Forward primer | Reverse primer |
|---|---|---|
| FN | TGAGCTGCACATGTCTTG | TCCTACGTGGTATGTCTTCC |
| GAPDH | AACTTTGGTATCGTGGAAGGAC | CAGTAGAGGCAGGGATGATGTT |
| CDH11 | TGGCAGCAAGTATCCAATGG | TTTGGTTACGTGGTAGGCAC |
| 18s | CATTCGAACGTCTGCCCTAT | CCTGCTGCCTTCCTTGGA |
| TNF | CAAAGCCCATGCACACTTCC | GGAAAGCCCCCTGTTTGAGT |
| IL6 | GGACCAAGACCATCCAATTC | ACCACAGTGAGGAATGTCCA |

Abbreviations: FN, Fibronectin; TNF, Tumor necrosis factor, IL, Interleukin

### Supplemental Table 4. Indices used for macroscopic and histological scoring of tissue specimen

#### A. Index for macroscopic tissue evaluation adapted from the simple endoscopic score for Crohn's disease (SES-CD)[1]

| | Surgical specimen (for each segment, ni/i/s) | | | |
|---|---|---|---|---|
| Variable | 0 | 1 | 2 | 3 |
| Size of ulcers | None | Small<br>(diameter 0.1-0.5 cm) | Medium<br>(diameter 0.5-2.0 cm) | Large<br>(diameter > 2.0 cm) |
| Ulcerated surface | None | < 10% | 10% – 30% | > 30% |
| Affected surface<br>- Ulcers<br>- Erosions<br>- Red discoloration<br>- Disappearance of the folds | Unaffected | < 50% | 50% – 75% | > 75% |
| Wall thickness<br>(without mesenteric fat) | None<br>(< 3 mm) | Mild<br>(3 – 5 mm) | Moderate<br>(> 5 – 7 mm) | Severe<br>(> 7 mm) |

#### B. Index for histological (inflammation and fibrosis) score of full thickness intestinal specimen

**(i) Inflammation score** (adapted from Geboes *et al.*[2])

| | **Structural (architectural change) (0.X)** | **Chronic inflammatory infiltrate (1.X)** | **LP Eosinophils (2A.X)** | **LP neutrophils (2B.X)** | **Epithelial neutrophils (3.X)** | **Crypt destruction (4.X)** | **Erosion or ulceration (5.X)** |
|---|---|---|---|---|---|---|---|
| **X.0** | No abnormality | No increase | No increase | No increase | No increase | None | None |
| **X.1** | Mild abnormality | Mild increase | Mild but unequivocal increase | Mild but unequivocal increase | <5% of crypts | Local excess of PMNs in part of a crypt | Recovering epithelium |
| **X.2** | Mild or moderate diffuse or multifocal abnormalities | Moderate increase | Moderate increase | Moderate increase | <50% of crypts | Marked attenuation | Probable erosion |

| | | | | | | | |
|---|---|---|---|---|---|---|---|
| **X.3** | Severe diffuse or multifocal abnormalities | Marked increase | Marked increase | Marked increase | >50% of crypts | Unequivocal crypt destruction | Unequivocal erosion |
| **X.4** | | | | | | | Ulcer or granulation tissue |
| The final score is the sum of the 7 subsocres and ranges from 0 to 22 | | | | | | | |

**(ii) Fibrosis (**adapted from Chiorean *et al.* [3])

| | | |
|---|---|---|
| 0 | No or minimal fibrosis | Limited to submucosa (<25% of thickness) |
| 1 | Mild | Submucosal fibrosis (25-50% of thickness) |
| 2 | Moderate | Submucosal fibrosis (>50% of thickness), septae into muscularis propria, muscular hyperplasia with preserved layers |
| 3 | Severe | Massive transmural fibrosis, effacement of normal layers |

**Supplemental Table 5. Cohort characteristics.**

**A.** Summary demographics

| Characteristic | CD Stricture | NL |
|---|---|---|
| | n = 13 | n = 7 |
| Age at time of surgery (Years), Median [IQR] | 42.0 [32.0] | 71.0 [14.0] |
| Age at diagnosis (Years), Median [IQR] | 24.0 [14] | 71.0 [13.5] |
| Gender, n (%) | | |
| Female | 8 (61.5) | 4 (57.1) |
| Male | 5 (38.5) | 3 (42.9) |
| Race, n (%) | | |
| American Indian | 0 (0) | 0 (0) |
| Asian | 0 (0) | 0 (0) |
| Black | 1 (7.7) | 3 (42.9) |
| Pacific Islander | 0 (0) | 0 (0) |
| White | 12 (92.3) | 4 (57.1) |
| Ethnicity, n (%) | | |
| Hispanic | 0 (0) | 0.0 |
| Non-Hispanic | 13 (100) | 7 (100) |
| Unknown | 0 (0) | 0.0 |
| BMI, Median [IQR] | 22.0 [7.97] | 24.0 [5.94] |
| Smoking Status, n (%) | | |
| Active | 2.0 (15.4) | 1.0 (14.3) |
| Former | 4 (30.8) | 2 (28.6) |
| Never | 7 (53.8) | 4 (57.1) |
| Anastomotic stricture, n (%) | | |
| No | 5 (38.5) | 7 (100.0) |
| Yes | 8 (61.5) | |
| Therapy at time of surgery, n (%) | | |
| None | 4 (30.8) | 7 (100.0) |
| 5-ASA | 2 (15.4) | |
| Anti-TNF | 6 (46.2) | |
| MTX | 1 (7.7) | |
| Perianal Disease, n (%) | | |
| No | 7 (53.8) | |
| Yes | 6 (46.2) | |
| Montreal Disease Classification (at any time during the disease course), n (%) | | |
| B2 | 6 (46.2) | |
| B2p | 4 (30.8) | |
| B3 | 1 (7.7) | |

| | |
|---|---|
| B3p | 2 (15.3) |

**B.** Patient level data corresponding to GEO upload

| Sample ID | Race | Gender | Diagnosis | Disease Location | Smoking Status | Perianal Disease | Therapy at time of Surgery | Macroscopic Inflammation Index* | | | |
|---|---|---|---|---|---|---|---|---|---|---|---|
| | | | | | | | | CDni | CDi | CDs | NL |
| CD2331 | White | Male | CD Stricture | Ileum | Never | No | None | 0 | 7 | 10 | NA |
| CD2375 | White | Female | CD Stricture | Ileum | Never | No | Anti-TNF | 0 | 4 | 11 | NA |
| CD2436 | White | Male | CD Stricture | Ileum | Former | No | MTX | 0 | 8 | 12 | NA |
| CD2440 | White | Male | CD Stricture | Ileum | Former | Yes | 5-ASA | 0 | 6 | 9 | NA |
| CD2459 | White | Male | CD Stricture | Ileum | Former | Yes | Anti-TNF | 0 | 5 | 10 | NA |
| CD2467 | White | Male | CD Stricture | Ileum | Never | Yes | None | 0 | 6 | 10 | NA |
| CD2469 | Black | Female | CD Stricture | Ileum | Never | No | Anti-TNF | 0 | 12 | 12 | NA |
| CD2484 | White | Female | CD Stricture | Ileum | Never | No | 5-ASA | 0 | 6 | 10 | NA |
| CD2577 | White | Female | CD Stricture | Ileum | Active | Yes | Anti-TNF | 0 | 8 | 12 | NA |
| CD2586 | White | Female | CD Stricture | Ileum | Active | No | None | 0 | 10 | 12 | NA |
| CD2587 | White | Female | CD Stricture | Ileum | Former | Yes | Anti-TNF | 0 | 3 | 6 | NA |
| CD2591 | White | Female | CD Stricture | Ileum | Never | No | None | 0 | 3 | 5 | NA |
| CD2598 | White | Female | CD Stricture | Ileum | Never | Yes | Anti-TNF | 0 | 8 | 12 | NA |
| NL2353 | Black | Male | NL | Colon | Never | No | None | NA | NA | NA | 0 |
| NL2402 | White | Male | NL | Colon | Former | No | None | NA | NA | NA | 0 |
| NL2449 | Black | Male | NL | Colon | Former | No | None | NA | NA | NA | 0 |
| NL2450 | White | Female | NL | Colon | Never | No | None | NA | NA | NA | 0 |
| NL2480 | White | Female | NL | Colon | Never | No | None | NA | NA | NA | 0 |
| NL2489 | White | Female | NL | Colon | Never | No | None | NA | NA | NA | 0 |
| NL2516 | Black | Female | NL | Colon | Active | No | None | NA | NA | NA | 0 |

*modified from the simple endoscopic score (SES-CD)

Abbreviations: CD, Crohn's disease; NL, normal; 5-ASA, 5-Aminosalicylates; TNF, Tumor necrosis factor; MTX, Methotrexate, NA; Not applicable

**C.** Histological scores of CD patient specimen used in the study

| Sample ID | Inflammation Score | | | Fibrosis score | | |
|---|---|---|---|---|---|---|
| | CDni | CDi | CDs | CDni | CDi | CDs |
| CD2331 | 0 | 15 | 11 | 0 | 2 | 3 |
| CD2375 | 0 | 2 | 16 | 0 | 1 | 2 |
| CD2436 | 0 | 13 | 11 | 0 | 1 | 2 |
| CD2440 | * | * | * | * | * | * |
| CD2459 | 0 | 5 | 8 | 0 | 2 | 0 |
| CD2467 | 0 | * | 7 | 0 | * | 2 |
| CD2469 | 0 | 4 | 13 | 0 | 0 | 3 |
| CD2484 | 0 | 3 | 9 | 0 | 0 | 3 |
| CD2577 | 0 | 17 | 13 | 0 | 1 | 1 |
| CD2586 | 0 | * | 13 | 0 | * | 2 |
| CD2587 | 0 | 10 | 13 | 0 | 2 | 2 |
| CD2591 | 0 | 8 | 9 | 0 | 2 | 3 |
| CD2598 | 0 | 7 | 5 | 0 | 2 | 2 |

*Insufficient tissue to evaluate

Abbreviations: CDs: strictured Crohn's disease; CDi: inflamed, non-strictured Crohn's disease; CDni: non-involved Crohn's disease

**Supplemental Table 6. Sequenced cell numbers for each compartment**

| Cell type | Total | CDs | CDi | CDni | NL |
|---|---|---|---|---|---|
| Epithelial cells | 125291 | 11010 | 35361 | 34258 | 44662 |
| Stromal cells | 96351 | 35185 | 16885 | 22137 | 22144 |
| Myeloid cells | 51163 | 17202 | 10523 | 10267 | 13171 |
| B cells | 78822 | 15388 | 22191 | 16655 | 24588 |
| T/NK cells | 57374 | 15264 | 16204 | 14637 | 11269 |

CDs: strictured Crohn's disease; CDi: inflamed, non-strictured Crohn's disease; CDni: non-inflamed Crohn's disease; NL: normal tissue

**Supplemental Table 7. Secretome analysis in primary human intestinal myofibroblast in response to cadherin-11 activation**

| Gene ID | Symbol | FC:CTRL ratio | Protein |
|---|---|---|---|
| 1937 | EEF1G | 0.243 | eukaryotic translation elongation factor 1 gamma |
| 5238 | PGM3 | 0.364 | phosphoglucomutase 3 |
| 10213 | PSMD14 | 0.437 | proteasome 26S subunit, non-ATPase 14 |
| 8417 | STX7 | 0.215 | syntaxin 7 |
| 26986 | PABPC1 | 0.236 | poly(A) binding protein cytoplasmic 1 |
| 378 | ARF4 | 0.259 | ADP ribosylation factor 4 |
| 8407 | TAGLN2 | 0.271 | transgelin 2 |
| 25802 | LMOD1 | 0.573 | leiomodin 1 |
| 3843 | IPO5 | 0.688 | importin 5 |
| 2547 | XRCC6 | 0.535 | X-ray repair cross complementing 6 |
| 10979 | FERMT2 | 0.38 | FERM domain containing kindlin 2 |
| 29952 | DPP7 | 0.852 | dipeptidyl peptidase 7 |
| 6218 | RPS17 | 0.246 | ribosomal protein S17 |
| 4904 | YBX1 | 0.208 | Y-box binding protein 1 |
| 3611 | ILK | 0.695 | integrin linked kinase |
| 5211 | PFKL | 0.355 | phosphofructokinase, liver type |
| 173 | AFM | 0.238 | afamin |
| 10417 | SPON2 | 0.183 | spondin 2 |
| 3956 | LGALS1 | 0.169 | galectin 1 |
| 10694 | CCT8 | 0.177 | chaperonin containing TCP1 subunit 8 |
| 71 | ACTG1 | 0.413 | actin gamma 1 |
| 5690 | PSMB2 | 0.197 | proteasome 20S subunit beta 2 |
| 28988 | DBNL | 0.82 | drebrin like |
| 81 | ACTN4 | 0.175 | actinin alpha 4 |
| 10383 | TUBB4B | 0.36 | tubulin beta 4B class IVb |
| 10097 | ACTR2 | 0.336 | actin related protein 2 |
| 10376 | TUBA1B | 0.25 | tubulin alpha 1b |
| 813 | CALU | 0.326 | calumenin |
| 4666 | NACA | 0.608 | nascent polypeptide associated complex subunit alpha |
| 1289 | COL5A1 | 0.428 | collagen type V alpha 1 chain |
| 2664 | GDI1 | 0.256 | GDP dissociation inhibitor 1 |
| 3178 | HNRNPA1 | 0.374 | heterogeneous nuclear ribonucleoprotein A1 |
| 159 | ADSS2 | 0.633 | adenylosuccinate synthase 2 |
| 4134 | MAP4 | 0.186 | microtubule associated protein 4 |
| 5660 | PSAP | 0.211 | prosaposin |
| 1660 | DHX9 | 0.285 | DExH-box helicase 9 |

| | | | |
|---|---|---|---|
| 2638 | GC | 0.198 | GC vitamin D binding protein |
| 171024 | SYNPO2 | 0.373 | synaptopodin 2 |
| 5518 | PPP2R1A | 0.301 | protein phosphatase 2 scaffold subunit Aalpha |
| 1278 | COL1A2 | 0.22 | collagen type I alpha 2 chain |
| 1072 | CFL1 | 0.304 | cofilin 1 |
| 4830 | NME1 | 0.238 | NME/NM23 nucleoside diphosphate kinase 1 |
| 7077 | TIMP2 | 0.308 | TIMP metallopeptidase inhibitor 2 |
| 396 | ARHGDIA | 0.211 | Rho GDP dissociation inhibitor alpha |
| 2665 | GDI2 | 0.277 | GDP dissociation inhibitor 2 |
| 10487 | CAP1 | 0.192 | cyclase associated actin cytoskeleton regulatory protein 1 |
| 5048 | PAFAH1B1 | 0.229 | platelet activating factor acetylhydrolase 1b regulatory subunit 1 |
| 3482 | IGF2R | 0.308 | insulin like growth factor 2 receptor |
| 7980 | TFPI2 | 0.983 | tissue factor pathway inhibitor 2 |
| 10575 | CCT4 | 0.307 | chaperonin containing TCP1 subunit 4 |
| 213 | ALB | 0.183 | albumin |
| 10611 | PDLIM5 | 0.912 | PDZ and LIM domain 5 |
| 10935 | PRDX3 | 0.473 | peroxiredoxin 3 |
| 11034 | DSTN | 0.206 | destrin, actin depolymerizing factor |
| 4478 | MSN | 0.241 | moesin |
| 140576 | S100A16 | 0.365 | S100 calcium binding protein A16 |
| 59 | ACTA2 | 0.549 | actin alpha 2, smooth muscle |
| 25932 | CLIC4 | 0.292 | chloride intracellular channel 4 |
| 55832 | CAND1 | 0.371 | cullin associated and neddylation dissociated 1 |
| 8826 | IQGAP1 | 0.279 | IQ motif containing GTPase activating protein 1 |
| 735 | C9 | 0.443 | complement C9 |
| 1264 | CNN1 | 0.408 | calponin 1 |
| 2331 | FMOD | 0.651 | fibromodulin |
| 22883 | CLSTN1 | 0.381 | calsyntenin 1 |
| 3094 | HINT1 | 0.403 | histidine triad nucleotide binding protein 1 |
| 3083 | HGFAC | 0.419 | HGF activator |
| 6125 | RPL5 | 0.498 | ribosomal protein L5 |
| 5859 | QARS1 | 1.04 | glutaminyl-tRNA synthetase 1 |
| 8411 | EEA1 | 0.422 | early endosome antigen 1 |
| 350 | APOH | 0.279 | apolipoprotein H |
| 2058 | EPRS1 | 0.248 | glutamyl-prolyl-tRNA synthetase 1 |
| 7171 | TPM4 | 0.351 | tropomyosin 4 |
| 720 | C4A | 0.205 | complement C4A (Rodgers blood group) |
| 203068 | TUBB | 0.413 | tubulin beta class I |
| 6251 | RSU1 | 0.429 | Ras suppressor protein 1 |
| 2274 | FHL2 | 0.472 | four and a half LIM domains 2 |
| 2318 | FLNC | 0.351 | filamin C |

| | | | |
|---|---|---|---|
| 22818 | COPZ1 | 0.457 | COPI coat complex subunit zeta 1 |
| 1627 | DBN1 | 0.361 | drebrin 1 |
| 11333 | PDAP1 | 0.849 | PDGFA associated protein 1 |
| 10540 | DCTN2 | 0.396 | dynactin subunit 2 |
| 3861 | KRT14 | 1.03 | keratin 14 |
| 7414 | VCL | 0.237 | vinculin |
| 1315 | COPB1 | 0.378 | COPI coat complex subunit beta 1 |
| 1266 | CNN3 | 0.367 | calponin 3 |
| 1973 | EIF4A1 | 0.303 | eukaryotic translation initiation factor 4A1 |
| 27044 | SND1 | 0.327 | staphylococcal nuclease and tudor domain containing 1 |
| 5213 | PFKM | 0.761 | phosphofructokinase, muscle |
| 7094 | TLN1 | 0.307 | talin 1 |
| 1933 | EEF1B2 | 0.314 | eukaryotic translation elongation factor 1 beta 2 |
| 4015 | LOX | 0.394 | lysyl oxidase |
| 10094 | ARPC3 | 0.301 | actin related protein 2/3 complex subunit 3 |
| 3045 | HBD | 0.473 | hemoglobin subunit delta |
| 3987 | LIMS1 | 0.703 | LIM zinc finger domain containing 1 |
| 3053 | SERPIND1 | 1.13 | serpin family D member 1 |
| 10095 | ARPC1B | 0.441 | actin related protein 2/3 complex subunit 1B |
| 8572 | PDLIM4 | 0.75 | PDZ and LIM domain 4 |
| 87 | ACTN1 | 0.294 | actinin alpha 1 |
| 56925 | LXN | 0.401 | latexin |
| 7335 | UBE2V1 | 0.344 | ubiquitin conjugating enzyme E2 V1 |
| 2317 | FLNB | 0.278 | filamin B |
| 10631 | POSTN | 0.266 | periostin |
| 5345 | SERPINF2 | 0.367 | serpin family F member 2 |
| 7791 | ZYX | 0.3 | zyxin |
| 2316 | FLNA | 0.308 | filamin A |
| 10272 | FSTL3 | 2.06 | follistatin like 3 |
| 1465 | CSRP1 | 0.692 | cysteine and glycine rich protein 1 |
| 2629 | GBA | 0.503 | glucosylceramidase beta |
| 55752 | SEPTIN11 | 0.364 | septin 11 |
| 10484 | SEC23A | 0.348 | SEC23 homolog A, COPII coat complex component |
| 1471 | CST3 | 0.564 | cystatin C |
| 4502 | MT2A | 0.651 | metallothionein 2A |
| 51637 | RTRAF | 0.707 | RNA transcription, translation and transport factor |
| 8615 | USO1 | 0.888 | USO1 vesicle transport factor |
| 4312 | MMP1 | 0.351 | matrix metallopeptidase 1 |
| 4314 | MMP3 | 0.555 | matrix metallopeptidase 3 |
| 3326 | HSP90AB1 | 0.249 | heat shock protein 90 alpha family class B member 1 |
| 351 | APP | 0.511 | amyloid beta precursor protein |

| | | | |
|---|---|---|---|
| 1265 | CNN2 | 0.637 | calponin 2 |
| 3490 | IGFBP7 | 0.361 | insulin like growth factor binding protein 7 |
| 10381 | TUBB3 | 0.5 | tubulin beta 3 class III |
| 7430 | EZR | 0.335 | ezrin |
| 7057 | THBS1 | 0.28 | thrombospondin 1 |
| 10516 | FBLN5 | 0.649 | fibulin 5 |
| 1277 | COL1A1 | 0.299 | collagen type I alpha 1 chain |
| 11167 | FSTL1 | 0.562 | follistatin like 1 |
| 1284 | COL4A2 | 0.533 | collagen type IV alpha 2 chain |
| 1938 | EEF2 | 0.379 | eukaryotic translation elongation factor 2 |
| 4026 | LPP | 0.463 | LIM domain containing preferred translocation partner in lipoma |
| 22872 | SEC31A | 0.364 | SEC31 homolog A, COPII coat complex component |
| 5358 | PLS3 | 0.343 | plastin 3 |
| 60 | ACTB | 0.381 | actin beta |
| 9260 | PDLIM7 | 0.442 | PDZ and LIM domain 7 |
| 800 | CALD1 | 0.501 | caldesmon 1 |
| 22820 | COPG1 | 0.394 | COPI coat complex subunit gamma 1 |
| 220 | ALDH1A3 | 0.587 | aldehyde dehydrogenase 1 family member A3 |
| 4860 | PNP | 0.554 | purine nucleoside phosphorylase |
| 7837 | PXDN | 0.666 | peroxidasin |
| 1462 | VCAN | 1.06 | versican |
| 23022 | PALLD | 0.816 | palladin, cytoskeletal associated protein |
| 1410 | CRYAB | 1.68 | crystallin alpha B |
| 9352 | TXNL1 | 0.597 | thioredoxin like 1 |
| 4237 | MFAP2 | 1.98 | microfibril associated protein 2 |
| 7076 | TIMP1 | 0.615 | TIMP metallopeptidase inhibitor 1 |
| 7130 | TNFAIP6 | 0.99 | TNF alpha induced protein 6 |
| 1490 | CCN2 | 3.61 | cellular communication network factor 2 |
| 2683 | B4GALT1 | 0.87 | beta-1,4-galactosyltransferase 1 |
| 7169 | TPM2 | 1.08 | tropomyosin 2 |
| 9590 | AKAP12 | 0.557 | A-kinase anchoring protein 12 |
| 4052 | LTBP1 | 2.43 | latent transforming growth factor beta binding protein 1 |
| 8482 | SEMA7A | 1.74 | semaphorin 7A (John Milton Hagen blood group) |
| 6876 | TAGLN | 0.472 | transgelin |
| 10468 | FST | 1.6 | follistatin |
| 1311 | COMP | 0.614 | cartilage oligomeric matrix protein |
| 1282 | COL4A1 | 1.57 | collagen type IV alpha 1 chain |
| 5236 | PGM1 | 0.822 | phosphoglucomutase 1 |
| 7168 | TPM1 | 1.49 | tropomyosin 1 |
| 1290 | COL5A2 | 0.72 | collagen type V alpha 2 chain |
| 2335 | FN1 | 0.707 | fibronectin 1 |

| | | | |
|---|---|---|---|
| 1809 | DPYSL3 | 0.8 | dihydropyrimidinase like 3 |
| 5069 | PAPPA | 2.07 | pappalysin 1 |
| 1466 | CSRP2 | 1 | cysteine and glycine rich protein 2 |
| 3624 | INHBA | 1.89 | inhibin subunit beta A |
| 5654 | HTRA1 | 0.944 | HtrA serine peptidase 1 |
| 29995 | LMCD1 | 1.97 | LIM and cysteine rich domains 1 |
| 6695 | SPOCK1 | 1.01 | SPARC (osteonectin), cwcv and kazal like domains proteoglycan 1 |
| 633 | BGN | 2.73 | biglycan |
| 3486 | IGFBP3 | 1.15 | insulin like growth factor binding protein 3 |
| 7045 | TGFBI | 1.17 | transforming growth factor beta induced |
| 9244 | CRLF1 | 1.32 | cytokine receptor like factor 1 |
| 4053 | LTBP2 | 1.35 | latent transforming growth factor beta binding protein 2 |
| 6678 | SPARC | 1.15 | secreted protein acidic and cysteine rich |
| 1000 | CDH2 | 2.09 | cadherin 2 |
| 1191 | CLU | 1.11 | clusterin |
| 1004 | CDH6 | 1.62 | cadherin 6 |
| 1314 | COPA | 1.23 | COPI coat complex subunit alpha |
| 5054 | SERPINE1 | 1.8 | serpin family E member 1 |
| 5270 | SERPINE2 | 2.48 | serpin family E member 2 |
| 4319 | MMP10 | 2.46 | matrix metallopeptidase 10 |

*$P < 0.05$

**Supplemental Table 8. Top 30 upregulated (A) and downregulated (B) genes in RNA sequencing analysis of HIMF after *CDH11* knockdown**

**A.**

| GeneID | Symbol | logFC | logCPM | PValue |
|---|---|---|---|---|
| 11009 | IL24 | 5.610 | 6.074 | 1.80589E-18 |
| 118430 | MUCL1 | 5.339 | 0.451 | 1.33564E-10 |
| 10578 | GNLY | 5.044 | -0.224 | 1.04409E-07 |
| 1437 | CSF2 | 4.966 | 2.887 | 3.81398E-17 |
| 100507056 | CCAT1 | 4.707 | -0.493 | 1.04935E-06 |
| 54210 | TREM1 | 4.672 | 1.179 | 4.8383E-11 |
| 80008 | TMEM156 | 4.549 | 2.196 | 1.29469E-14 |
| 118421 | LINC00161 | 4.409 | 0.793 | 2.90624E-10 |
| 1440 | CSF3 | 4.207 | 1.426 | 8.89048E-12 |
| 338322 | NLRP10 | 4.169 | -0.513 | 4.17651E-06 |
| 4312 | MMP1 | 4.029 | 9.953 | 1.84714E-28 |
| 597 | BCL2A1 | 3.945 | 1.764 | 4.56691E-12 |
| 6364 | CCL20 | 3.896 | 2.340 | 6.38593E-15 |
| 5055 | SERPINB2 | 3.786 | 4.011 | 7.52377E-20 |
| 55117 | SLC6A15 | 3.687 | 3.308 | 4.77801E-17 |
| 1880 | GPR183 | 3.597 | -0.651 | 3.06445E-05 |
| 5743 | PTGS2 | 3.591 | 9.478 | 4.4024E-30 |
| 3553 | IL1B | 3.574 | 6.522 | 4.65538E-19 |
| 3552 | IL1A | 3.553 | 1.726 | 5.95718E-11 |
| 2494 | NR5A2 | 3.457 | 0.597 | 5.24148E-08 |
| 90865 | IL33 | 3.415 | 7.974 | 1.70227E-11 |
| 8988 | HSPB3 | 3.260 | -0.304 | 6.62322E-06 |
| 7056 | THBD | 3.219 | 1.484 | 7.59747E-10 |
| 345651 | ACTBL2 | 3.170 | -0.370 | 2.92128E-05 |
| 26298 | EHF | 3.160 | 1.190 | 4.14084E-09 |
| 441389 | LINC01239 | 3.154 | 0.140 | 1.22241E-06 |
| 4773 | NFATC2 | 3.076 | 0.381 | 1.91808E-06 |
| 114 | ADCY8 | 3.039 | 1.195 | 1.14603E-08 |
| 103752588 | PACERR | 2.945 | 0.080 | 1.95112E-05 |
| 245972 | ATP6V0D2 | 2.927 | 0.341 | 3.27933E-06 |

**B.**

| GeneID | Symbol | logFC | logCPM | PValue |
|---|---|---|---|---|
| 118427 | OLFM3 | -5.666 | -0.169 | 3.65624E-08 |
| 1009 | CDH11 | -5.126 | 8.046 | 4.30361E-34 |
| 170689 | ADAMTS15 | -4.672 | 2.630 | 2.80975E-16 |
| 23209 | MLC1 | -4.328 | 0.345 | 1.45324E-08 |
| 11095 | ADAMTS8 | -4.056 | 4.753 | 5.55714E-23 |
| 6750 | SST | -3.904 | 3.318 | 1.51499E-16 |
| 29951 | PDZRN4 | -3.902 | 3.767 | 3.00257E-19 |
| 125 | ADH1B | -3.852 | 5.403 | 2.23418E-25 |
| 55959 | SULF2 | -3.659 | 7.698 | 6.01603E-30 |
| 3479 | IGF1 | -3.417 | 2.169 | 7.84581E-12 |
| 6387 | CXCL12 | -3.403 | 8.377 | 6.69472E-30 |
| 9201 | DCLK1 | -3.359 | 2.217 | 9.74734E-11 |
| 1E+08 | PRORY | -3.291 | 1.434 | 6.15079E-10 |
| 1E+08 | NPY4R2 | -3.248 | 0.509 | 4.16322E-06 |
| 9547 | CXCL14 | -3.247 | 6.070 | 1.61788E-25 |
| 730 | C7 | -3.182 | 0.641 | 3.16814E-07 |
| 3038 | HAS3 | -3.164 | 3.052 | 5.46923E-16 |
| 23090 | ZNF423 | -3.141 | 1.367 | 5.77134E-09 |
| 1591 | CYP24A1 | -3.133 | 1.195 | 3.53537E-08 |
| 9934 | P2RY14 | -3.058 | 0.355 | 3.31527E-06 |
| 643650 | LINC00842 | -2.981 | 2.427 | 1.0845E-11 |
| 5136 | PDE1A | -2.939 | 3.610 | 8.67812E-17 |
| 7373 | COL14A1 | -2.913 | 8.998 | 3.15374E-24 |
| 653145 | ANXA8 | -2.912 | 1.625 | 9.07693E-09 |
| 3122 | HLA-DRA | -2.891 | -0.125 | 8.94641E-05 |
| 6355 | CCL8 | -2.878 | 1.380 | 1.90631E-08 |
| 8630 | HSD17B6 | -2.863 | 3.216 | 5.60405E-15 |
| 11249 | NXPH2 | -2.859 | 3.203 | 1.26964E-14 |
| 57125 | PLXDC1 | -2.839 | 1.259 | 3.29753E-08 |
| 23671 | TMEFF2 | -2.744 | 1.658 | 2.18909E-08 |

**Supplemental Table 9.** Correlation of *CDH11* with core extracellular matrix (ECM) genes

| **GeneID** | **Symbol** | **corn** | **GeneName** |
|---|---|---|---|
| 7373 | COL14A1 | 0.567 | collagen, type XIV, alpha 1 |
| 4147 | MATN2 | 0.504 | matrilin 2 |
| 25878 | MXRA5 | 0.496 | matrix-remodelling associated 5 |
| 6586 | SLIT3 | 0.437 | slit homolog 3 (Drosophila) |
| 4239 | MFAP4 | 0.387 | microfibrillar-associated protein 4 |
| 10418 | SPON1 | 0.384 | spondin 1, extracellular matrix protein |
| 127435 | PODN | 0.381 | podocan |
| 83716 | CRISPLD2 | 0.374 | cysteine-rich secretory protein LCCL domain containing 2 |
| 131873 | COL6A6 | 0.363 | collagen, type VI, alpha 6 |
| 2202 | EFEMP1 | 0.352 | EGF-containing fibulin-like extracellular matrix protein 1 |
| 7148 | TNXB | 0.34 | tenascin XB |
| 80781 | COL18A1 | 0.334 | collagen, type XVIII, alpha 1 |
| 284654 | RSPO1 | 0.333 | R-spondin homolog (Xenopus laevis) |
| 340419 | RSPO2 | 0.332 | R-spondin 2 homolog (Xenopus laevis) |
| 2192 | FBLN1 | 0.329 | fibulin 1 |
| 25992 | SNED1 | 0.303 | sushi, nidogen and EGF-like domains 1 |
| 84570 | COL25A1 | 0.301 | collagen, type XXV, alpha 1 |
| 5549 | PRELP | 0.301 | proline/arginine-rich end leucine-rich repeat protein |
| 1298 | COL9A2 | 0.299 | collagen, type IX, alpha 2 |
| 2201 | FBN2 | 0.298 | fibrillin 2 |
| 129804 | FBLN7 | 0.294 | fibulin 7 |
| 129080 | EMID1 | 0.285 | EMI domain containing 1 |
| 5118 | PCOLCE | 0.275 | procollagen C-endopeptidase enhancer |
| 5212 | VIT | 0.27 | vitrin |
| 79875 | THSD4 | 0.259 | thrombospondin, type I, domain containing 4 |
| 2199 | FBLN2 | 0.258 | fibulin 2 |
| 79883 | PODNL1 | 0.256 | podocan-like 1 |
| 64856 | VWA1 | 0.256 | von Willebrand factor A domain containing 1 |
| 1277 | COL1A1 | 0.253 | collagen, type I, alpha 1 |
| 10516 | FBLN5 | 0.248 | fibulin 5 |
| 5649 | RELN | 0.247 | reelin |
| 1311 | COMP | 0.243 | cartilage oligomeric matrix protein |
| 1634 | DCN | 0.241 | decorin |
| 255743 | NPNT | 0.239 | nephronectin |
| 3339 | HSPG2 | 0.232 | heparan sulfate proteoglycan 2 |
| 1842 | ECM2 | 0.224 | extracellular matrix protein 2, female organ and adipocyte specific |
| 3908 | LAMA2 | 0.223 | laminin, alpha 2 |
| 1289 | COL5A1 | 0.222 | collagen, type V, alpha 1 |

| | | | |
|---|---|---|---|
| 2331 | FMOD | 0.217 | fibromodulin |
| 7448 | VTN | 0.207 | vitronectin |
| 1287 | COL4A5 | 0.201 | collagen, type IV, alpha 5 |
| 1306 | COL15A1 | 0.195 | collagen, type XV, alpha 1 |
| 10085 | EDIL3 | 0.191 | EGF-like repeats and discoidin I-like domains 3 |
| 633 | BGN | 0.189 | biglycan |
| 84624 | FNDC1 | 0.188 | fibronectin type III domain containing 1 |
| 83690 | CRISPLD1 | 0.186 | cysteine-rich secretory protein LCCL domain containing 1 |
| 4060 | LUM | 0.185 | lumican |
| 8425 | LTBP4 | 0.184 | latent transforming growth factor beta binding protein 4 |
| 285313 | IGSF10 | 0.182 | immunoglobulin superfamily, member 10 |
| 3909 | LAMA3 | 0.182 | laminin, alpha 3 |
| 4811 | NID1 | 0.179 | nidogen 1 |
| 1293 | COL6A3 | 0.177 | collagen, type VI, alpha 3 |
| 3910 | LAMA4 | 0.176 | laminin, alpha 4 |
| 1303 | COL12A1 | 0.174 | collagen, type XII, alpha 1 |
| 2006 | ELN | 0.172 | elastin |
| 4256 | MGP | 0.17 | matrix Gla protein |
| 3486 | IGFBP3 | 0.17 | insulin-like growth factor binding protein 3 |
| 1462 | VCAN | 0.164 | versican |
| 7059 | THBS3 | 0.163 | thrombospondin 3 |
| 2200 | FBN1 | 0.161 | fibrillin 1 |
| 1281 | COL3A1 | 0.152 | collagen, type III, alpha 1 |
| 84034 | EMILIN2 | 0.149 | elastin microfibril interfacer 2 |
| 1307 | COL16A1 | 0.147 | collagen, type XVI, alpha 1 |
| 2621 | GAS6 | 0.146 | growth arrest-specific 6 |
| 1296 | COL8A2 | 0.143 | collagen, type VIII, alpha 2 |
| 3488 | IGFBP5 | 0.143 | insulin-like growth factor binding protein 5 |
| 10631 | POSTN | 0.142 | periostin, osteoblast specific factor |
| 4053 | LTBP2 | 0.137 | latent transforming growth factor beta binding protein 2 |
| 83872 | HMCN1 | 0.13 | hemicentin 1 |
| 3912 | LAMB1 | 0.121 | laminin, beta 1 |
| 1290 | COL5A2 | 0.119 | collagen, type V, alpha 2 |
| 79987 | SVEP1 | 0.117 | sushi, von Willebrand factor type A, EGF and pentraxin domain containing 1 |
| 1301 | COL11A1 | 0.116 | collagen, type XI, alpha 1 |
| 30008 | EFEMP2 | 0.114 | EGF-containing fibulin-like extracellular matrix protein 2 |
| 165 | AEBP1 | 0.113 | AE binding protein 1 |
| 1302 | COL11A2 | 0.107 | collagen, type XI, alpha 2 |
| 1101 | CHAD | 0.107 | chondroadherin |
| 3487 | IGFBP4 | 0.106 | insulin-like growth factor binding protein 4 |
| 25987 | TSKU | 0.104 | tsukushi small leucine rich proteoglycan homolog (Xenopus laevis) |

| | | | |
|---|---|---|---|
| 3915 | LAMC1 | 0.103 | laminin, gamma 1 (formerly LAMB2) |
| 9353 | SLIT2 | 0.102 | slit homolog 2 (Drosophila) |
| 1291 | COL6A1 | 0.101 | collagen, type VI, alpha 1 |
| 5552 | SRGN | 0.1 | serglycin |
| 1278 | COL1A2 | 0.099 | collagen, type I, alpha 2 |
| 6678 | SPARC | 0.097 | secreted protein, acidic, cysteine-rich (osteonectin) |
| 59277 | NTN4 | 0.095 | netrin 4 |
| 168667 | BMPER | 0.095 | BMP binding endothelial regulator |
| 4237 | MFAP2 | 0.094 | microfibrillar-associated protein 2 |
| 4054 | LTBP3 | 0.093 | latent transforming growth factor beta binding protein 3 |
| 25890 | ABI3BP | 0.083 | ABI family, member 3 (NESH) binding protein |
| 11117 | EMILIN1 | 0.081 | elastin microfibril interfacer 1 |
| 342035 | GLDN | 0.08 | gliomedin |
| 145864 | HAPLN3 | 0.072 | hyaluronan and proteoglycan link protein 3 |
| 22932 | POMZP3 | 0.072 | POM (POM121 homolog, rat) and ZP3 fusion |
| 4240 | MFGE8 | 0.064 | milk fat globule-EGF factor 8 protein |
| 1292 | COL6A2 | 0.062 | collagen, type VI, alpha 2 |
| 9423 | NTN1 | 0.055 | netrin 1 |
| 3911 | LAMA5 | 0.054 | laminin, alpha 5 |
| 6695 | SPOCK1 | 0.053 | sparc/osteonectin, cwcv and kazal-like domains proteoglycan (testican) 1 |
| 1893 | ECM1 | 0.051 | extracellular matrix protein 1 |
| 1404 | HAPLN1 | 0.047 | hyaluronan and proteoglycan link protein 1 |
| 7045 | TGFBI | 0.046 | transforming growth factor, beta-induced, 68kDa |
| 8292 | COLQ | 0.045 | collagen-like tail subunit (single strand of homotrimer) of asymmetric acetylcholinesterase |
| 3913 | LAMB2 | 0.028 | laminin, beta 2 (laminin S) |
| 50509 | COL5A3 | 0.022 | collagen, type V, alpha 3 |
| 2335 | FN1 | 0.022 | fibronectin 1 |
| 7784 | ZP3 | 0.02 | zona pellucida glycoprotein 3 (sperm receptor) |
| 79174 | CRELD2 | 0.019 | cysteine-rich with EGF-like domains 2 |
| 80144 | FRAS1 | 0.018 | Fraser syndrome 1 |
| 4013 | VWA5A | 0.016 | von Willebrand factor A domain containing 5A |
| 1285 | COL4A3 | 0.015 | collagen, type IV, alpha 3 (Goodpasture antigen) |
| 3489 | IGFBP6 | 0.014 | insulin-like growth factor binding protein 6 |
| 89932 | PAPLN | 0.014 | papilin, proteoglycan-like sulfated glycoprotein |
| 3371 | TNC | 0.005 | tenascin C |
| 1286 | COL4A4 | 0.004 | collagen, type IV, alpha 4 |
| 4238 | MFAP3 | 0.002 | microfibrillar-associated protein 3 |

**Supplemental Table 10. KEGG pathways enriched with *CDH11* knockdown in HIMF. (A) Pathways enriched with upregulated genes in *CDH11* knockdown. (B) Pathways enriched with downregulated genes in *CHD11* knockdown.**

**A.**

| ID | Description | GeneRatio | BgRatio | pvalue | p.adjust | Count |
|---|---|---|---|---|---|---|
| hsa04060 | Cytokine-cytokine receptor interaction | 23/97 | 295/8163 | 2.93394E-13 | 6.19061E-11 | 23 |
| hsa05164 | Influenza A | 17/97 | 171/8163 | 1.14718E-11 | 1.21027E-09 | 17 |
| hsa04657 | IL-17 signaling pathway | 13/97 | 94/8163 | 5.65709E-11 | 3.97882E-09 | 13 |
| hsa05171 | Coronavirus disease - COVID-19 | 18/97 | 232/8163 | 1.73269E-10 | 9.13992E-09 | 18 |
| hsa05323 | Rheumatoid arthritis | 12/97 | 93/8163 | 7.4706E-10 | 3.15259E-08 | 12 |
| hsa04668 | TNF signaling pathway | 12/97 | 112/8163 | 6.58543E-09 | 2.31588E-07 | 12 |
| hsa05162 | Measles | 12/97 | 139/8163 | 7.63981E-08 | 2.30286E-06 | 12 |
| hsa04061 | Viral protein interaction with cytokine and cytokine receptor | 10/97 | 100/8163 | 2.51217E-07 | 6.62586E-06 | 10 |
| hsa05160 | Hepatitis C | 11/97 | 157/8163 | 2.24691E-06 | 5.26775E-05 | 11 |
| hsa04623 | Cytosolic DNA-sensing pathway | 7/97 | 63/8163 | 8.64341E-06 | 0.000182376 | 7 |
| hsa04622 | RIG-I-like receptor signaling pathway | 7/97 | 70/8163 | 1.7502E-05 | 0.00033572 | 7 |
| hsa04620 | Toll-like receptor signaling pathway | 8/97 | 104/8163 | 2.99922E-05 | 0.000527363 | 8 |
| hsa05417 | Lipid and atherosclerosis | 11/97 | 215/8163 | 4.46646E-05 | 0.000724941 | 11 |
| hsa05202 | Transcriptional misregulation in cancer | 10/97 | 193/8163 | 9.11581E-05 | 0.001373883 | 10 |
| hsa04630 | JAK-STAT signaling pathway | 9/97 | 162/8163 | 0.000122761 | 0.001726841 | 9 |
| hsa04640 | Hematopoietic cell lineage | 7/97 | 99/8163 | 0.000164256 | 0.002166122 | 7 |
| hsa04933 | AGE-RAGE signaling pathway in diabetic complications | 7/97 | 100/8163 | 0.000174923 | 0.002171108 | 7 |
| hsa05133 | Pertussis | 6/97 | 76/8163 | 0.000269471 | 0.003158802 | 6 |
| hsa05144 | Malaria | 5/97 | 50/8163 | 0.000296434 | 0.003291976 | 5 |
| hsa04621 | NOD-like receptor signaling pathway | 9/97 | 184/8163 | 0.000320281 | 0.003378963 | 9 |
| hsa05167 | Kaposi sarcoma-associated herpesvirus infection | 9/97 | 194/8163 | 0.000472673 | 0.004749238 | 9 |
| hsa05169 | Epstein-Barr virus infection | 9/97 | 202/8163 | 0.000633767 | 0.006002716 | 9 |
| hsa05161 | Hepatitis B | 8/97 | 162/8163 | 0.000654324 | 0.006002716 | 8 |
| hsa05142 | Chagas disease | 6/97 | 102/8163 | 0.001295913 | 0.011393233 | 6 |
| hsa05163 | Human cytomegalovirus infection | 9/97 | 225/8163 | 0.001362164 | 0.011496661 | 9 |

| | | | | | | |
|---|---|---|---|---|---|---|
| hsa04625 | C-type lectin receptor signaling pathway | 6/97 | 104/8163 | 0.00143284 | 0.011596412 | 6 |
| hsa05332 | Graft-versus-host disease | 4/97 | 42/8163 | 0.001483901 | 0.011596412 | 4 |
| hsa04940 | Type I diabetes mellitus | 4/97 | 43/8163 | 0.001621346 | 0.012218004 | 4 |
| hsa04062 | Chemokine signaling pathway | 8/97 | 192/8163 | 0.001958426 | 0.014249235 | 8 |
| hsa04010 | MAPK signaling pathway | 10/97 | 294/8163 | 0.002481734 | 0.017454865 | 10 |
| hsa04913 | Ovarian steroidogenesis | 4/97 | 51/8163 | 0.003054036 | 0.020787149 | 4 |
| hsa05134 | Legionellosis | 4/97 | 57/8163 | 0.004573066 | 0.030153653 | 4 |
| hsa05166 | Human T-cell leukemia virus 1 infection | 8/97 | 222/8163 | 0.004776405 | 0.030540042 | 8 |
| hsa04350 | TGF-beta signaling pathway | 5/97 | 94/8163 | 0.005095608 | 0.031622746 | 5 |
| hsa05146 | Amoebiasis | 5/97 | 102/8163 | 0.007179477 | 0.042748752 | 5 |
| hsa05321 | Inflammatory bowel disease | 4/97 | 65/8163 | 0.007293626 | 0.042748752 | 4 |
| hsa04064 | NF-kappa B signaling pathway | 5/97 | 104/8163 | 0.007781454 | 0.044375321 | 5 |

**B.**

| ID | Description | GeneRatio | BgRatio | pvalue | p.adjust | Count |
|---|---|---|---|---|---|---|
| hsa00600 | Sphingolipid metabolism | 4/93 | 53/8163 | 0.003018671 | 0.049670856 | 4 |
| hsa04151 | PI3K-Akt signaling pathway | 13/93 | 354/8163 | 0.000171673 | 0.007768214 | 13 |
| hsa04512 | ECM-receptor interaction | 7/93 | 88/8163 | 5.94737E-05 | 0.003588245 | 7 |
| hsa04514 | Cell adhesion molecules | 7/93 | 157/8163 | 0.002021757 | 0.04544035 | 7 |
| hsa04510 | Focal adhesion | 9/93 | 201/8163 | 0.000447851 | 0.013510158 | 9 |
| hsa05410 | Hypertrophic cardiomyopathy | 9/93 | 90/8163 | 7.23196E-07 | 0.000113616 | 9 |
| hsa05414 | Dilated cardiomyopathy | 9/93 | 96/8163 | 1.25543E-06 | 0.000113616 | 9 |
| hsa05412 | Arrhythmogenic right ventricular cardiomyopathy | 6/93 | 77/8163 | 0.00023004 | 0.008327448 | 6 |
| hsa04640 | Hematopoietic cell lineage | 6/93 | 99/8163 | 0.00088981 | 0.023007934 | 6 |
| hsa05205 | Proteoglycans in cancer | 8/93 | 205/8163 | 0.002259465 | 0.04544035 | 8 |
| hsa04610 | Complement and coagulation cascades | 5/93 | 85/8163 | 0.002757492 | 0.049670856 | 5 |
